# A *FLOWERING LOCUS T* ortholog is associated with photoperiod-insensitive flowering in hemp (*Cannabis sativa* L.)

**DOI:** 10.1101/2023.04.21.537862

**Authors:** Caroline A. Dowling, Jiaqi Shi, Jacob A. Toth, Michael A. Quade, Lawrence B. Smart, Paul F. McCabe, Rainer Melzer, Susanne Schilling

## Abstract

Hemp (*Cannabis sativa* L.) is an extraordinarily versatile crop, with applications ranging from medicinal compounds to seed oil and fibre products. *Cannabis sativa* is a short-day plant, and its flowering is tightly controlled by photoperiod. However, substantial genetic variation exists for photoperiod sensitivity in *C. sativa,* and photoperiod-insensitive (“autoflower”) cultivars are available.

Using a bi-parental mapping population and bulked segregant analysis, we identified *Autoflower2*, a 0.5 Mbp locus significantly associated with photoperiod-insensitive flowering in hemp. *Autoflower2* contains an ortholog of the central flowering time regulator *FLOWERING LOCUS T* (*FT*) from *Arabidopsis thaliana* which we termed *CsFT1*. Extensive sequence divergence between alleles of *CsFT1* was identified between photoperiod-sensitive and insensitive cultivars of *C. sativa*, including a duplication of *CsFT1* and sequence differences especially in introns. Genotyping of several mapping populations and a diversity panel confirmed a strong correlation between *CsFT1* alleles and photoperiod response as well as affirming that at least two independent loci for this agriculturally important trait, *Autoflower1* and *Autoflower2*, exist in the *C. sativa* gene pool.

This study reveals the multiple independent origins of photoperiod insensitivity in *C. sativa,* supporting the likelihood of a complex domestication history in this species. By integrating the genetic relaxation of photoperiod sensitivity into novel *C. sativa* cultivars, expansion to higher latitudes will be permitted, thus allowing the full potential of this versatile crop to be reached.

## Introduction

The precise timing of flowering is crucial to reproductive success (Blümel *et al*., 2015). Flowering time is controlled by an intricate network of environmental and endogenous cues that converge to signal the transition from vegetative to reproductive plant growth (Cao *et al*., 2021). Domestication permitted expansion of the world’s major crops to diverse ecological conditions, and this phenomenon depended on key alterations in flowering time genes (Gaudinier and Blackman, 2020; Hill and Li, 2016). As a yield-related trait, fine-tuning flowering time is a major goal of crop improvement efforts, specifically to develop new crop varieties adapted to different latitudes (Cao *et al*., 2021).

*Cannabis sativa* L. is one of the earliest known cultivated plants, with archaeological records implying human use at least 8,000 years before present (Kovalchuk *et al*., 2020; Long *et al*., 2017). A multipurpose crop, the applications of *C. sativa* are significant, spanning biofuel, textiles, building materials and paper (Schilling *et al*., 2021). Additionally, as a fast-growing annual plant, *C. sativa* has the potential to aid carbon sequestration efforts (Finnan and Styles, 2013). However, *C. sativa* is best known for its medicinal and intoxicating applications, as glandular trichomes on female *C. sativa* flowers produce high-value, psychoactive compounds called cannabinoids (Livingston *et al*., 2020; Melzer *et al*., 2022). One of these cannabinoids, cannabidiol (CBD), is non-intoxicating with anti-cancer and anti-inflammatory effects, currently used to treat conditions like epilepsy and multiple sclerosis (Schilling *et al*., 2021). Another, Δ^9^-tetrahydrocannabinol (THC) is an intoxicating cannabinoid used for recreational and therapeutic purposes. The regulatory landscape surrounding *C. sativa* is complex, rapidly evolving, and country-specific. *Cannabis sativa* is legally classified as either hemp (below 0.3% THC) or marijuana (above 0.3% THC).

Breeders of hemp have selected lines with flowering time appropriate for crop applications or market class. Connections between fibre quality and flowering time are recognised, with early flowering cultivars typically favoured for seed production whereas late flowering cultivars are favoured for fibre production (Salentijn *et al*., 2019). As a short-day species, flowering in *C. sativa* is affected by light conditions with optimal, cultivar-specific photoperiods usually in the range of ∼12-15 hours (Lisson *et al*., 2000; Zhang *et al*., 2021). Therefore, developing new cultivars adapted to a wide geographic range and suited to the final market class of the crop can be facilitated by a comprehensive understanding of the genetic underpinnings of flowering (Dowling *et al*., 2021).

Alongside *Humulus lupulus*, *C. sativa* was one of the first plants for which the critical role of the photoperiod in determining flowering was described (Tournois 1912, cited in Heslop-Harrison, 1956; Kobayashi and Weigel, 2007). Substantial flowering time variation exists in *C. sativa*, with observable differences between cultivars (Stack *et al*., 2021). Furthermore, day-neutral or “autoflowering” cultivars exist that are photoperiod insensitive and can flower under continuous light. Photoperiod insensitivity provides opportunities for *C. sativa* cultivation at higher latitudes or in controlled environments under long day conditions and these cultivars typically mature faster and are smaller in height (Stack *et al*., 2021). Previous research demonstrated that in the hemp cultivar ‘FINOLA’, the cultivar central to this study, flowering occurs independent of day length (Callaway and Laakkonen, 1996; Schilling *et al*., 2023) (https://finola.fi/, last accessed on 20 Mar., 2023).

Here, we present the identification of *Autoflower2*, a locus contributing to photoperiod-insensitive flowering in ‘FINOLA’. We applied a modified QTL-seq approach combining bulked segregant analysis and whole-genome sequencing (WGS) (Itoh *et al*., 2019). Using this approach we identified a candidate gene located at the end of chromosome (chr) 8, an ortholog of *FLOWERING LOCUS T* (*FT*): *CsFT1*. Sequence analysis and in-depth genotyping reveal a tandem duplication of *CsFT1*. We hypothesise that *CsFT1* is involved in photoperiod-insensitive flowering in ‘FINOLA’. Genotyping of a diverse panel of accessions confirms multiple independent origins of photoperiod insensitivity in *C. sativa*.

## Methods

### Plant materials, growth conditions and phenotyping

Hemp (*Cannabis sativa* L.) parent plants were selected that differ in their sensitivity to photoperiod. ‘FINOLA’ is a photoperiod-insensitive grain cultivar, and ‘Felina 32’ is a photoperiod-sensitive dual-purpose grain/fibre cultivar. A cross between a monoecious ‘Felina 32’ individual (male flowers removed) and a male ‘FINOLA’ individual was used to generate a ‘Felina 32’ × ‘FINOLA’ F_1_ population. To generate the F_2_, a single male F_1_ plant was used to pollinate four F_1_ female plants (sibling cross). The parent plants and F_1_ individuals that were used to generate the F_2_ population were cultivated using a speed breeding protocol previously described (Schilling *et al*., 2023).

The ‘Felina 32’ × ‘FINOLA’ F_2_ population consisted of 245 individuals with offspring from all four F_1_ female individuals. The cultivation of the F_2_ population took place under natural light conditions in a glasshouse in Dublin, Ireland, between June 1^st^ and September 25^th^ 2020. Alongside the F_2_, parental plants ‘FINOLA’ (n = 30) and ‘Felina 32’ (n = 11), as well as F_1_ plants (n = 47), were grown to permit phenotypic comparison with the F_2_ population. The F_3_ population (n=250) was generated by random cross pollination of the F_2_ plants, which was approximately 31.5% monoecious. The F_3_ population was cultivated in a glasshouse in Dublin, Ireland between June 1^st^ and September 28^th^ 2021.

A commercial fertilizer (Miracle-Gro, NPK 24-8-16 and Miracle-Gro Magnesium Salts, https://scottsmiraclegro.com/) was applied weekly to plants in all conditions to prevent nutrient deficiency. Temperature, humidity, and day length were recorded throughout both glasshouse trials (Figure S1). In 2022, ‘FINOLA’ (n=38) and ‘Felina 32’ (n=16) individuals were grown under continuous light at 26°C in 50-cell deep trays in glasshouses at Cornell AgriTech (Geneva, NY) and flowering times recorded.

Seven populations from the Cornell hemp breeding program were evaluated in 50-cell deep trays under continuous light at Cornell AgriTech (Geneva, NY) to score for photoperiod sensitivity. ‘Picolo’ (Verve Seed Solutions, Saskatoon, Saskatchewan, Canada) is an early maturing Canadian grain-type cultivar closely related to ‘FINOLA’ (Carlson *et al*., 2021) and previously shown to flower under continuous light (Toth *et al*., 2022), while GVA-H-20-1179 (hereafter called 20-1179) is a late-flowering Cornell breeding line derived from a Chinese landrace. The F_1_ population (GVA-H-21-1130) was developed by crossing a single female 20-1179 with four male ‘Picolo’. Subsequently, an F_2_ population (GVA-H-22-1103) was generated by crossing one F_1_ female with a sibling F_1_ male. The F_1_ backcross population (GVA-H-22-1104) was generated by crossing a single female 20-1179 (late flowering parent) with a single F_1_ male (GVA-H-21-1130). Numerous individuals from each of the lines in this pedigree (20-1179, n=24; ‘Picolo’, n=16; F_1_ population GVA-H-21-1130, n=44; F_2_ population GVA-H-22-1103, n=92; and F_1_ backcross population GVA-H-22-1104, n=67) were grown under continuous light to score for photoperiod sensitivity. These lines, except for the F_1_ backcross, were also grown outdoors under organic field conditions in Geneva, NY. Seeds of ‘Picolo’, 20-1179 and the F_2_ population were sown in 50-cell deep trays on May 9, 2022. DNA marker screening of seedlings of ‘Picolo’ and 20-1179 was conducted using CSP-2 (Toth, 2022) to identify and remove males. The female F_1_ parent used to generate the F_2_ population was propagated from cuttings on June 9, 2022. Female seedlings of the parents (‘Picolo’, n=11; 20-1179, n=16), rooted cuttings of the F_1_ female parent (n=24), and dioecious seedlings of the F_2_ population (n=339) were transplanted to the field on July 8, 2022. Rating for flowering time was done weekly with female flowering time defined as the stage when at least one pair of stigma was visible. Male flowering was defined as when individual male flowers larger than 2 mm were visible.

An F_1_ population (GVA-H-21-1139) was developed by crossing a single female of the photoperiod-insensitive cultivar ‘Le Crème’ (Ventura Seed Company, Camarillo, CA, USA) homozygous for *Autoflower1* (Toth *et al*., 2022) with the same four male ‘Picolo’ plants used to generate the 20-1179 × ‘Picolo’ F_1_ population (GVA-H-21-1130) described above. ‘Le Crème’ × ‘Picolo’ F_2_ families were produced by crossing female F_1_ plants with multiple male F_1_ plants, with each half-sibling family (GVA-H-22-1176, GVA-H-22-1177, GVA-H-22-1178, GVA-H-22-1179, GVA-H-22-1180) representing seed collected from a single female parent. These F_2_ families (n=263) were also grown under continuous light to score for photoperiod sensitivity. Screening of individuals for flowering was conducted three times a week, using the same criteria described above. The CSP-2 sex marker was used to determine the sex of non-flowering individuals under continuous light (Toth, 2022).

For the *C. sativa* diversity panel, leaves and seeds (n=4-8 per accession) were obtained from the Cornell hemp program. ‘Russian Auto CBD’ seeds were sourced from Native Canna, Austria (https://www.nativcanna.at/shop/hanfsamen/russian-auto-cbd/).

### Nucleic acid isolation, Illumina sequencing and quality control

For whole genome sequencing (WGS), DNA was extracted from young leaves of parents, ‘Felina 32’ × ‘FINOLA’ F_1_, female F_2_ and F_3_ plants and a female ‘FINOLA’ using the DNeasy Plant Mini kit (Qiagen, Germany). For the Cornell hemp breeding program populations DNA was isolated as previously described (Toth *et al*., 2022). For the *C. sativa* diversity panel, DNA was isolated from leaves and seeds using the E-Z 384 Plant DNA HT kit (Omega Bio-tek, GA, USA).

For WGS DNA was quantified with Qubit dsDNA HS Assay Kit (Invitrogen, USA, catalogue number Q32851), permitting precise combination of individual DNA samples into bulks in equal ratios. Bulks for early and late flowering of 5 and 10 individuals each were created for ‘Felina 32’ × ‘FINOLA’ F_2_ and F_3_ generation, respectively, and individual DNA samples were mixed prior to sequencing.

All WGS samples were sequenced generating 150 bp paired-end reads (Illumina Novaseq, Novogene Europe, Cambridge, UK) with a target coverage of 30X for the two parent plants and the female ‘FINOLA’, 10X coverage for the three F_1_ individuals, 10X coverage for the eight F_2_ bulks (four early and four late flowering, each 5 individuals) and 20X coverage for the two F_3_ bulks (one early and one late flowering, each 10 individuals) (approximately 2X per individual in each bulk).

Raw sequencing reads were processed using the Galaxy platform (Afgan *et al*., 2018). Quality of reads was inspected (FastQC v. 0.11.8, https://www.bioinformatics.babraham.ac.uk/projects/fastqc, last accessed on 7 Mar., 2023) and low-quality reads and any adapter contamination were removed (Trimmomatic v. 0.38.1, ILLUMINACLIP:TruSeq3-PE.fa:2:30:10:8:TRUE; LEADING:10; TRAILING:10; SLIDINGWINDOW:5:15; MINLEN:100; AVGQUAL:20) (Bolger *et al*., 2014).

### Bulked segregant analysis using QTL-seq

To identify loci associated with early flowering, a ‘modified QTL-seq’ method, designed for highly heterozygous genomes, was utilised (Itoh *et al*., 2019). The pipeline was used with default parameters, using the photoperiod-sensitive parent ‘Felina 32’ as “parent 1” and the *C. sativa* ‘CBDRx’ reference genome (‘CBDRx’-cs10 v2) (Grassa *et al*., 2021; https://www.ncbi.nlm.nih.gov/assembly/GCF_900626175.2/, last accessed on 10 Jan., 2023). For scripts and key parameters see supplementary methods. An additional QTL-seq pipeline (Sugihara *et al*., 2022) was used to repeat the mapping using the publicly available ‘FINOLA’ genome (Laverty *et al*., 2019, https://www.ncbi.nlm.nih.gov/assembly/GCA_003417725.2, last accessed on 2 Jul., 2020) as the reference.

### RNA-seq analysis

Publicly available *C. sativa* RNA-seq data were obtained from NCBI and analysed (Table S1). Read quality was examined (FastQC v. 0.11.8) and filtering removed low-quality reads and adapter contamination, with Trimmomatic (v. 0.38.1) settings the same as genomic sequencing data except the following: ILLUMINACLIP adjusted for the sequencing technology used; Minimum read length cut-off was reduced according to sequencer read length: MINLEN:40 for 75-150 bp reads, or MINLEN:20 if reads below 40 bp (Bolger et al., 2014). Reads were mapped to the ‘CBDRx’ v2 reference genome using HISAT2 v2.2.1 (Kim *et al*., 2015). RSeQC Infer Experiment tool was used to determine strandedness of mapped datasets (Wang *et al*., 2012). StringTie was used to assemble transcripts and calculate gene expression in transcripts per million (TPM) (Pertea *et al*., 2015). Morpheus (https://software.broadinstitute.org/morpheus, last accessed on 24 Mar., 2023) was used to generate heatmaps from TPM values (hierarchical clustering, Euclidean distance).

### Analysis of genomic and protein sequences

Coverage calculations and plots of the genomic locus of *CsFT1* were created by mapping reads to ‘CBDRx’ and the ‘FINOLA’ genome (GCA_003417725.2) using BWA-MEM (Galaxy version v0.7.17.1), samtools depth (v1.10) and R (v3.6.0.1) (Li *et al*., 2009; Li and Durbin, 2010; R Core Team, 2022) (Supplementary methods).

For sequence analysis of *CsFT1* (LOC115700781, XM_030628422.1), *CsELF9* (LOC115699158) and Cs*AGO5a* (LOC115698998) and *CsAGO5b* (LOC115700443), mRNA and genomic sequences were obtained from NCBI. The genomic sequences of the ‘FINOLA’ and ‘Felina 32’ alleles of these genes were generated by mapping the sequencing data to the reference genome (GCF_900626175.2). To obtain the genomic location of *CsFT1* in the ‘FINOLA’ genome, tblastn was used with *CsFT1* as a query (Boratyn *et al*., 2013). To obtain the genomic sequence of the *CsFT1a* and *CsFT1b* copies, short reads of the ‘FINOLA’ parent were mapped to the ‘FINOLA’ genome (GCA_003417725.2) and the consensus sequence obtained in IGV. Jalview was used to translate cDNA to protein and both Jalview and EMBOSS Water (https://www.ebi.ac.uk/Tools/psa/emboss_water/, last accessed on 19 Jan., 2023) were used to conduct pairwise similarity comparisons of coding and protein sequences (Madeira *et al*., 2022). Protein domains of candidate genes were verified with HMMER, NCBI CDD and InterPro (Lu *et al*., 2020; Paysan-Lafosse *et al*., 2023; Potter *et al*., 2018).

### Phylogenetic analysis of *PEBP*-like genes

For comparative sequence analysis of the *PEBP* gene family, protein sequences were obtained from TAIR (The Arabidopsis Information Resource, www.arabidopsis.org, last accessed on 5 Sept., 2022) for *A. thaliana* and from literature (Venail *et al*., 2022) and downloaded from Phytozome (Goodstein *et al*., 2012; Ouyang *et al*., 2007) for *Oryza sativa* (Table S2). Protein sequences of *C. sativa* PEPB-like genes in ‘CBDRx’ were identified using BLAST with *A. thaliana* PEPB proteins as query. For all identified putative *C. sativa* PEPB-like proteins, the presence of the PEBP domain was verified using the NCBI conserved domain database (Lu *et al*., 2020). Sequence alignments were created using MAFFT (Katoh *et al*., 2019) and visualised in Jalview (Katoh *et al*., 2019; Waterhouse *et al*., 2009). ALISTAT v.1.3 was used to mask residues in the sequence alignment with a completeness score of > 0.5 (Wong *et al*., 2014). A maximum likelihood phylogenetic tree was constructed using IQ-TREE including branch supports from Ultrafast bootstraps and a Shi-modaira–Hasegawa approximate likelihood ratio test (SH-aLRT) test (1000 replicates each) (Trifinopoulos *et al*., 2016). FigTree was employed for tree visualisation (http://tree.bio.ed.ac.uk/software/figtree/, last accessed on 3 Apr., 2023).

### Genotyping of the *CsFT1* locus

To develop markers to assist breeding, a PCR Allele Competitive Extension (PACE™, 3CR Bioscience, UK) SNP genotyping assay was employed to target a G/A SNP difference between ‘Felina 32’ and ‘FINOLA’ alleles of *CsFT1* (A in ‘Felina 32’, G in both *CsFT1a* and *CsFT1b* in ‘FINOLA’ exon 4, NC_044379.1:64421558). Two specific differentially tagged forward primers and a single common reverse primer were designed (Table S3). The PACE reaction was conducted as previously described (Toth *et al*., 2020), using the above *CsFT1* primers and *Autoflower1* PACE primers (Toth *et al*., 2022). CFX Maestro v1.1 software (Bio-Rad, Hercules, CA, USA) was used for allelic discrimination analysis.

To screen individuals for copy number variation of *CsFT1*, quantitative PCR using fast SYBR™ Green kit (Thermofisher, USA) and ViiA 7 Real-Time PCR machine (Thermofisher, USA) was employed. DNA was isolated as described previously for WGS. Primers complementary to exon 1 of all *CsFT1* alleles (Table S3) were used for qPCR using genomic DNA as template. Results were normalised using the 2^-ddCt^ method with ‘Felina 32’ as a calibrator given the apparent single *CsFT1* copy in this cultivar. Furthermore, *CsLEAFY* (*CsLFY*, LOC115695615), a known single copy gene (Sayou *et al*., 2014), was used as the internal control for each sample to correct for variability in the DNA concentration of samples (Table S3).

## Results

### A mapping population from a cross between photoperiod-sensitive and photoperiod-insensitive hemp cultivars segregated for flowering time

To investigate photoperiod sensitivity in hemp, plants of the cultivars ‘FINOLA’ and ‘Felina 32’ were grown in the glasshouse under natural light conditions (long days, June to September) in two successive growing seasons (Figure 1a, b). Female individuals of the dioecious cultivar ‘FINOLA’ flowered significantly earlier than individuals of the monoecious cultivar ‘Felina 32’ under those conditions (2020: P = 1.1e-0.6, 2021: P = 2.22e-16, Mann–Whitney U-test, Figure 1c, S2a), indicating that ‘FINOLA’ and ‘Felina 32’ have a different photoperiodic response. This is in line with previous descriptions of ‘FINOLA’ being a photoperiod-insensitive hemp cultivar where flowering is independent of day length (Schilling *et al*., 2023). To investigate the genetic control of photoperiod-insensitive flowering time in hemp, a ‘Felina 32’ plant was cross-pollinated with a male ‘FINOLA’ individual (Figure 1d). The resulting ‘Felina 32’ × ‘FINOLA’ F_1_ individuals were exclusively dioecious with an intermediate flowering time between, and statistically different from, both parents (*P* < 0.05, Figure 1e, S2). Plants of the ‘Felina 32’ × ‘FINOLA’ F_2_ displayed wide variation in flowering time from 42 to 90 d after sowing (Figure 1e, S2). An F_3_ population generated by random intercrossing of F_2_ individuals similarly displayed a wide range of flowering times (42-80 d after sowing) (Figure 1e). In all populations, male individuals consistently flowered before female and monoecious individuals (Figure S2b).

**Figure 1.**
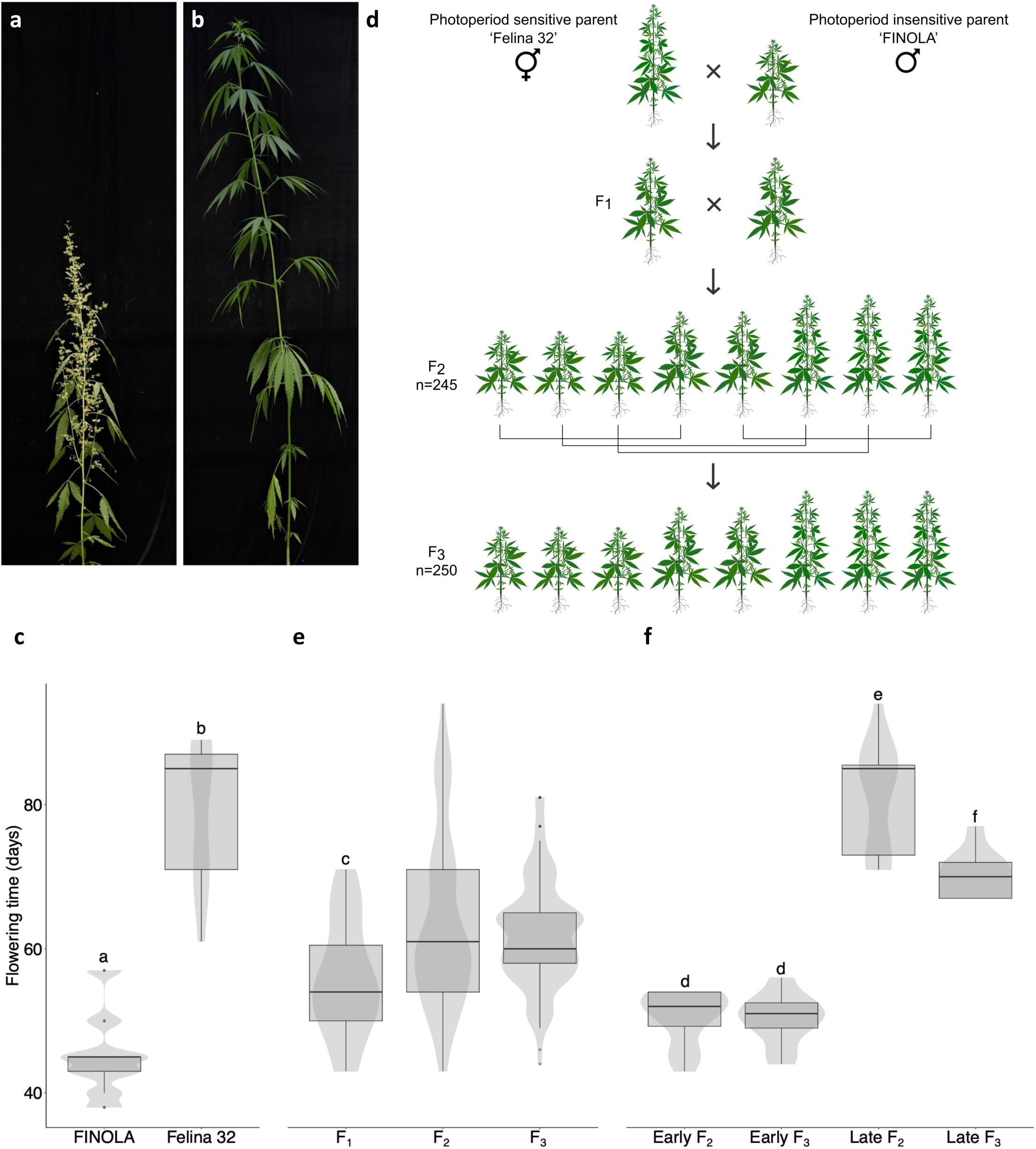
A hemp population segregating for photoperiod response. ‘FINOLA’ (a) and ‘Felina 32’ (b) differ in photoperiod-response. Under glasshouse conditions with natural light, ‘FINOLA’ flowered significantly earlier than ‘Felina 32’ (c). A mapping population was generated by crossing a monoecious ‘Felina 32’ individual (female parent) with a male ‘FINOLA’ plant and subsequently conducting sibling crosses in F_1_ and F_2_ (d). Flowering times of female individuals from the ‘Felina 32’ × ‘FINOLA’ F_1_, F_2_ and F_3_ generation are displayed (e). Bulks of female individuals with early vs. later flowering times were created from F_2_ and F_3_ individuals to be used in QTL-seq analysis (f). In (c), (e) and (f) significance levels are indicated by letters (*P* < 0.05, Mann–Whitney *U*-test).

### Identification of a locus controlling photoperiod-insensitive flowering in *C. sativa* via bulked segregant analysis and QTL-seq

To identify the genomic locus controlling photoperiod-insensitive flowering in ‘FINOLA’, a bulked segregant method was used (Itoh *et al*., 2019). For both ‘Felina 32’ × ‘FINOLA’ F_2_ and F_3_ populations we generated bulks for early and later flowering with 20 individuals each (Figure 1f). Using the F_2_ sequencing data we detected several QTLs for flowering time (Figure 2a, b, Table S4, S5, S6). When QTL-seq was repeated using the F_3_ sequencing data, only two QTLs were found: one QTL on chr08 (NC_044379.1:64.1-64.6 Mb, 0.5 Mb) present also in the F_2_-based analysis, and a QTL on chr03 (NC_044372.1:26.5-26.8 Mb, size 0.3Mb) which was not present in the F_2_-based analysis (Figure 2a, c, Table S4, S7). The QTL on chr08 was reduced in size in the F_3_-based analysis when compared to the F_2_-based analysis: from 2.4 Mb containing 159 genes, to 0.5 Mb containing 38 protein-coding genes (Figure 2, Table S4, S8). As the QTL on chr08 was the only one that was identified in both the ‘Felina 32’ × ‘FINOLA’ F_2_ and F_3_ population, we focussed our subsequent analyses on this locus, terming it *Autoflower2* in line with the previously identified *Autoflower1* locus for “autoflowering” in a high cannabinoid *C. sativa* cultivar (Toth *et al*., 2022).

**Figure 2.**
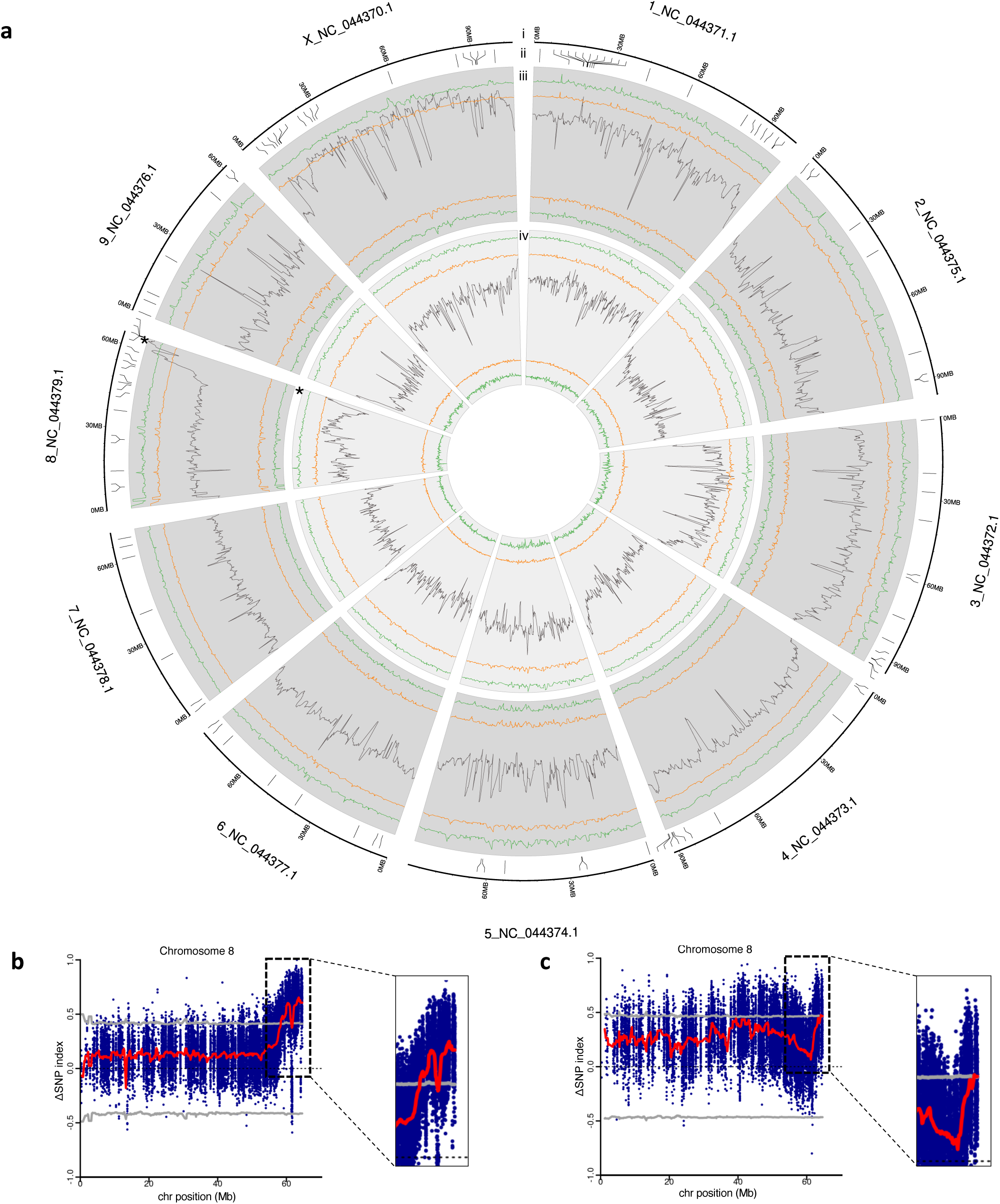
Bulked segregant analysis reveals a genetic locus associated with photoperiod-insensitive flowering in hemp. A circos plot (a) denoting the reference genome chromosomes (i, ‘CBDRx’, GCF_900626175.2), candidate genes for flowering time control in *C. sativa* (ii, Table S6), and sliding window (1-Mb interval, 100-kb increment) averages of ΔSNP indices derived from QTL-seq analysis of the ‘Felina 32’ × ‘FINOLA’ F_2_ (iii) and F_3_ (iv) generations. In (iii) and (iv) orange and green lines denote 95% and 99% confidence intervals, respectively. Several significant QTLs were detected (Table S4), but only the QTL at the end of chr08 was identified in both ‘Felina 32’ × ‘FINOLA’ F_2_ and F_3_ population (asterisk). ΔSNP-index plots of chr08 for the F_2_ (b) and F_3_ (c) population show the location of *Autoflower2* at the end of chr08. The sliding window averages of ΔSNP indices are depicted as red lines, individual SNP indices are displayed as blue dots. Circos plot was created using shinyCircos (Yu *et al*., 2018).

Subsequently, we reanalysed the sequencing data using the previously mentioned QTL-seq pipeline this time using the publicly available ‘FINOLA’ genome (Laverty *et al*., 2019) as the reference. Here, none of the assembled chromosomes presented a significant QTL. Therefore, 9 unassembled contigs containing the locus syntenic to the QTL on chr08 of the reference genome were analysed. Six out of nine analysed contigs showed significant hits (Figure S3).

### *Autoflower2* contains four candidate genes for flowering time control in *C. sativa*

To identify possible candidate genes for photoperiod-insensitive flowering in hemp we analysed the QTL *Autoflower2* in more detail. Among the genes located within the confidence interval of *Autoflower2* (Table S8), four have high similarity to known flowering time regulators from *A. thaliana*: *FLOWERING LOCUS T* (*FT*), *EARLY FLOWERING9* (*ELF9*) and two that are similar to *ARGONAUTE5* (*AGO5*). We therefore named the corresponding *C. sativa* genes *CsFT1* (LOC115700781), *CsELF9* (LOC115699158), Cs*AGO5a* (LOC115698998) and *CsAGO5b* (LOC115700443).

To assess whether these candidate genes are transcribed, and therefore potentially functional, the expression of all genes in *Autoflower2* was analysed using publicly available RNA-seq datasets (Figure 3). Both *AGO5a* and *AGO5b* were very weakly expressed in trichomes, flowers, leaves, bast fibres, stems, roots and mixed tissues. *CsELF9* was highly expressed in all analysed tissues, whereas high *CsFT1* expression was observed in flowers, bast fibres, stems, roots and seeds, with low expression in some leaf samples.

**Figure 3.**
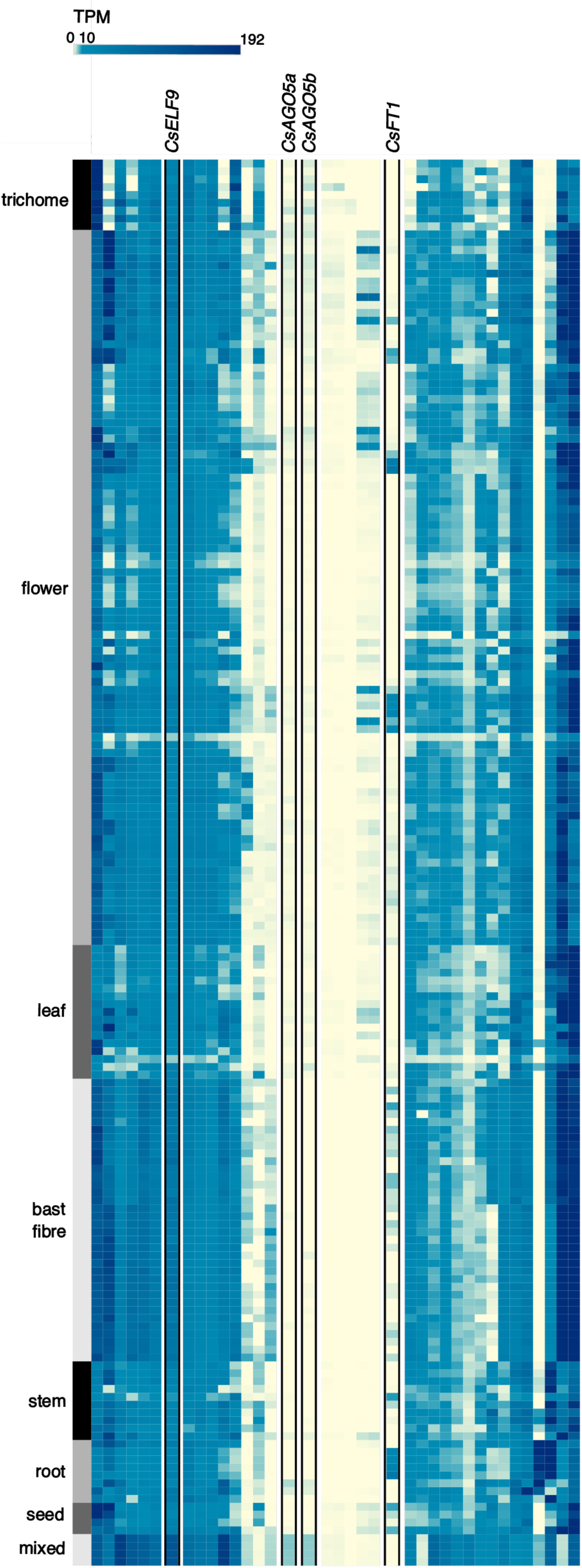
Expression patterns of all genes contained in *Autoflower2*. A heatmap displaying the expression of all genes in *Autoflower2*. Expression data in TPM were generated using publicly available RNA-seq datasets (Table S1).

We next compared predicted protein and coding sequences of *CsAGO5a*, *CsAGO5b*, *CsELF9* and *CsFT1* of the ‘FINOLA’ and ‘Felina 32’ cultivars. CsAGO5a and CsELF9 were highly similar between ‘FINOLA’ and ‘Felina 32’ (>98%, Table S9) while CsAGO5b sequence similarity was slightly lower (about 95%, Table S9). Differences in the CsELF9 sequence were due to a single amino acid substitution and an indel in conserved domains (Figure S4). The predicted sequences of CsAGO5a and CsAGO5b differed in a number of single amino acid substitutions within conserved domains (Figure S5).

Sequence comparisons of CsFT1 between ‘FINOLA’ and ‘Felina 32’ showed similar differences in predicted protein sequences (about 95%, Table S9). In addition to minor differences in the CDS, there seemed to be large structural differences between the alleles, including between introns, prompting a more detailed sequence analysis.

### *CsFT1* alleles show variation in structure and sequence between hemp cultivars

The *Autoflower2* locus contains 38 genes, including *CsFT1* and other candidate genes for flowering time control (Figure 4a). In the reference genome ‘CBDRx’ (Grassa *et al*., 2021), a single copy of *CsFT1* is present (Figure 4b). To compare the genomic architecture of *CsFT1* alleles, we located *CsFT1* in the published ‘FINOLA’ genome (Laverty et al., 2019, GCA_003417725.2). Here, *CsFT1* is located on a contig (QKVJ02000894.1:190-200kb) with two copies of *CsFT1* separated by ∼3.2 kb (Figure 4c, S6a). An alignment-based dot plot indicated that the duplication was restricted to only the *FT*-like genes, and there was large structural variation especially in the introns of *CsFT1* (Figure S6b). We designated the tandem duplicated copies found in ‘FINOLA’ *CsFT1a* and *CsFT1b* and annotated exons of both genes according to similarity to *CsFT1* in ‘CBDRx’.

**Figure 4.**
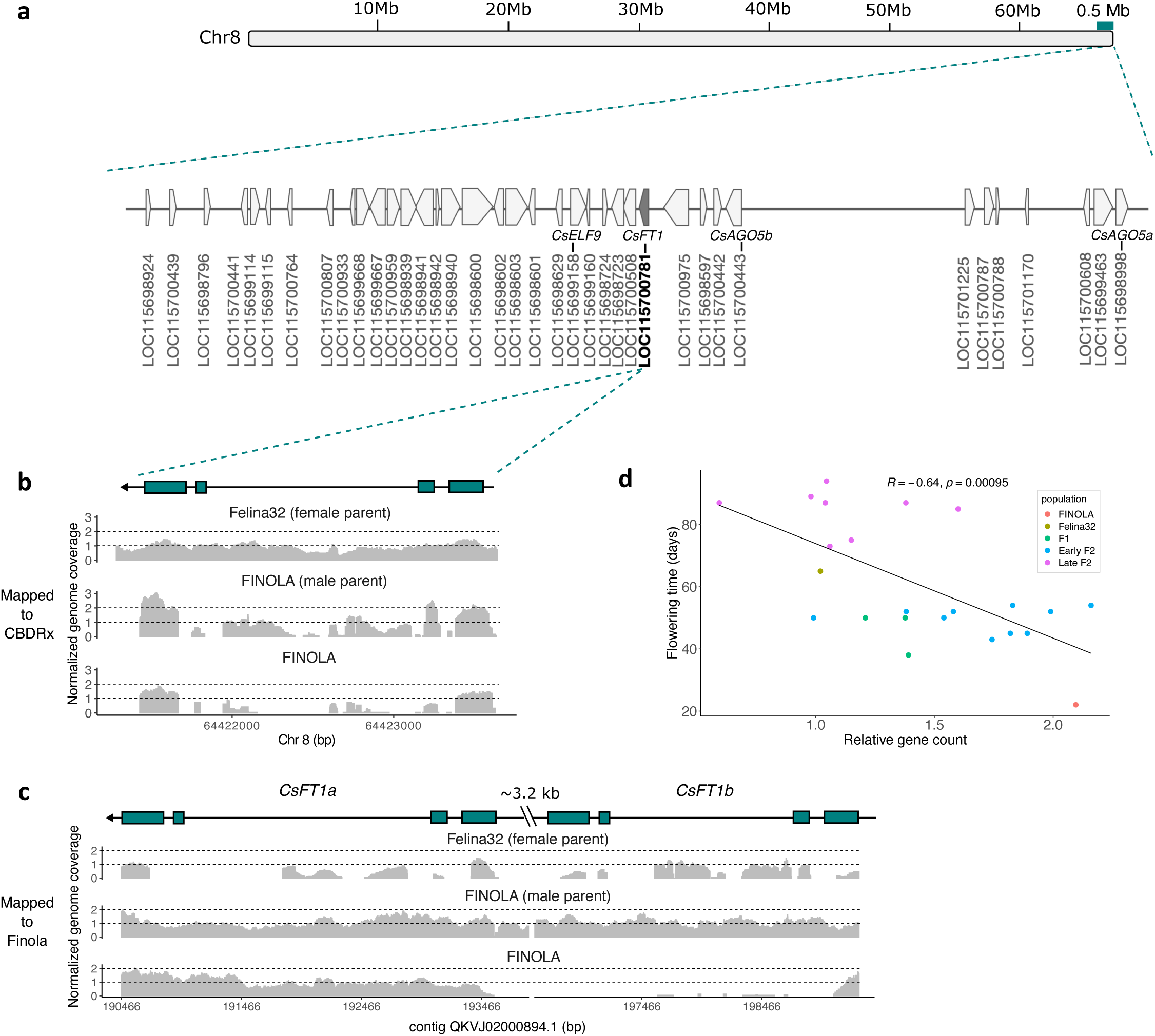
Structural variant in a candidate gene in *Autoflower2*. *Autoflower2* is located at the end of chr08 and contains 38 protein coding genes (a). Normalised coverage at *CsFT1* was calculated from short-read whole genome sequencing data mapped to the *C. sativa* reference genome ‘CBDRx’ (Grassa *et al*., 2021) (b) and the publicly available ‘FINOLA’ genome (Laverty *et al*., 2019) (c). Coverage was calculated from reads derived from sequencing the photoperiod-sensitive parent of the ‘Felina 32’ × ‘FINOLA’ population, ‘Felina 32’, the photoperiod-insensitive parent ‘FINOLA’ (male parent), and an additional female ‘FINOLA’ individual. (d) Flowering time and relative copy number of *CsFT1* are negatively correlated (Pearson’s coefficient= -0.64, P < 0.05).

The notion that *CsFT1* is tandem duplicated in the photoperiod-insensitive ‘FINOLA’ but only present as a single copy in ‘Felina 32’ was further supported by coverage analysis derived from WGS short-read data (Figure 4b). Sequence reads generated from the photoperiod-sensitive ‘Felina 32’ parent covered the genomic locus of *CsFT1* at an expected coverage of 1X when normalised to the whole genome (normalised coverage = 1X in exons, Figure 4b). Conversely, reads generated from the ‘FINOLA’ parent resulted in around twice the expected coverage in exons (1.9X). In introns, which are expected to have lower similarity between cultivars, coverage was slightly lower than in exons for ‘Felina 32’ (0.8X in introns vs. 1X in exons). The difference between coverage in exons and introns was much more pronounced for reads from ‘FINOLA’ (0.6X in introns vs. 1.9X in exons), indicating intronic sequence divergence between ‘FINOLA’ and the reference genome (Figure 4b).

Subsequently, we repeated the coverage analysis by mapping sequence reads to the ‘FINOLA’ genome (Laverty *et al*., 2019). For the ‘FINOLA’ parent the coverage was continuous between exons and introns and comparable to the average genome-wide coverage (approx. 1X over both *CsFT1a* and *CsFT1b*, Figure 4c). In contrast, reads generated from the ‘Felina 32’ parent poorly mapped to both exons and introns (0.6X over both copies, Figure 4c).

Additionally, we sequenced the genome of a second ‘FINOLA’ individual (short-read WGS). For this individual, when mapped to ‘CBDRx’, the coverage at *CsFT1* was similar to the ‘Felina 32’ sample in exons, but very low in introns (1.2X and 0.3X, respectively, Figure 4b). When mapped to the ‘FINOLA’ genome (Laverty *et al*., 2019), the coverage of *CsFT1a* was as expected while coverage over *CsFT1b* was low overall (1X and 0.3X, respectively, Figure 4c). This might indicate that not all ‘FINOLA’ individuals have a tandem duplication of *CsFT1*. To verify the apparent duplication of *CsFT1* in ‘FINOLA’, we used qPCR with genomic DNA as template, calculating a normalised relative gene count using the single copy gene *CsLFY* (Sayou *et al*., 2014). Relative gene count of *CsFT1* was higher in early flowering ‘Felina 32’ × ‘FINOLA’ F_2_ and ‘FINOLA’ individuals than in later flowering F_2_ and ‘Felina 32’ individuals and was observed to be negatively correlated with flowering time (R=-0.64, P = 0.00095, Figure 4d).

After detecting the copy number variation of *CsFT1* we wanted to understand sequence variation between the alleles and the putative gene duplicates. *CsFT1* found in the photoperiod-sensitive ‘Felina 32’ was identical to the reference genome. In contrast, the sequences of CsFT1a and CsFT1b differ from the reference sequence (95 and 98%, respectively, Table S9), and also from one another (96%).

CsFT1a is less similar to CsFT1 (8 amino acids different, 95% similarity) than CsFT1b (3 amino acids different, 98% similarity, Table S9), with all amino acid changes located in the PEBP domain (Figure 5a). One substitution in both CsFT1a and CsFT1b is located in the binding interface for 14-3-3 proteins (position 91, Figure 5a). In *A. thaliana*, FT has the same amino acid at this position as CsFT1a and CsFT1b, and this residue has not been demonstrated to be critical for protein interactions (Pieper *et al*., 2021; Taoka *et al*., 2011).

**Figure 5.**
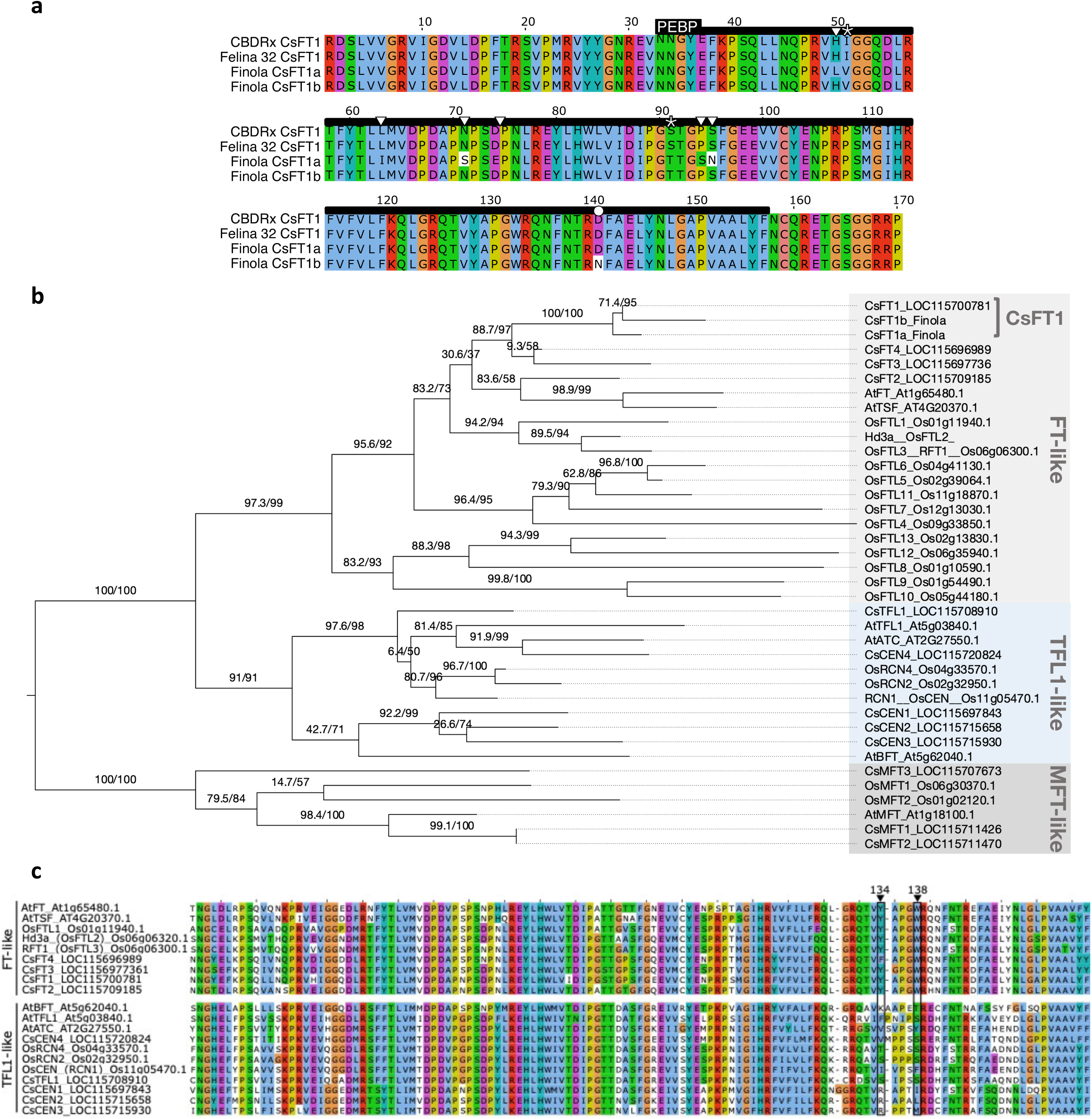
*CsFT1* is a PEBP-like protein and a likely flowering promoter in *C. sativa*. An alignment of CsFT1 protein predictions based on the *C. sativa* reference genome (‘CBDRx’) as well as sequencing data of ‘Felina32’ and ‘FINOLA’ (AA19-188, range identified by NCBI CDD as PEBP domain) shows sequence differences between alleles (a). CsFT1a and CsFT1b differ in eight and three amino acids respectively when compared to CsFT1 ‘Felina 32’ (asterisks) whereas CsFT1a and CsFT1b differ in seven amino acids from one another (triangles). A phylogenetic tree of PEPB-like proteins from *A. thaliana*, *O. sativa* and *C. sativa* shows three main clades: FT, MFT and TFL (b). An alignment of the PEPB domain region of PEBP-like proteins from *A. thaliana*, *O. sativa* and *C. sativa* highlights highly conserved and variable sites (c). Amino acid positions 134 and 138 (black boxes) are known to be important for distinguishing flowering promoters from repressors (Wickland and Hanzawa, 2015). For better visualisation only *O. sativa* PEBP of known function (Table S2) are shown.

Furthermore, located in the second intron of *CsFT1a* is a ∼1000 bp long section with similarity to multiple other loci throughout the genome (Figure S7a, b). The repetitive nature of the sequence was evidence for a mobile genetic element, though extended sequence similarities to known DNA or RNA transposable elements could not be detected. However, 5’ and 3’ ends of the sequence are similar to the consensus sequences of helitrons, a highly variable type of DNA transposable elements without highly conserved sequence or structural elements (Figure S7c) (Krasileva, 2019).

### CsFT1 is a putative flowering promoter

CsFT1 has high similarity to FT from *A. thaliana*. In order to analyse the phylogenetic relationship between the proteins, a maximum likelihood tree was constructed on the basis of a protein alignment, including all *C. sativa*, *A. thaliana* and *O. sativa* orthologs of FT, MOTHER OF FT (MFT) and TERMINAL FLOWER 1 (TFL) (Table S2). PEPB-domain proteins from *O. sativa* and *A. thaliana* were previously identified to be involved in flowering time control in both model species (Kardailsky *et al*., 1999; Kobayashi *et al*., 1999; Nakagawa *et al*., 2002; Song *et al*., 2018; Tamaki *et al*., 2007; Yoo *et al*., 2004). Within the phylogenetic tree, we observed three conserved and previously reported clades: FT-like, MFT-like and TFL-like (Figure 5b). The FT clade containing FT and HEADING DATE 3A, the *O. sativa* FT ortholog, is well supported (SH-LRT/UF bootstrap 97.3/99). CsFT1 is positioned within this group, confirming a close phylogenetic relationship. Alongside CsFT1 three additional *C. sativa* proteins fall within the FT clade, hence termed CsFT2, CsFT3 and CsFT4. The tree further confirms that CsFT1a and CsFT1b are closely related to CsFT1 (100/100). The paralogs CsFT1, 3 and 4 are contained in one well supported clade (88.7/97), while FT, TSF and CsFT2 appear to be more closely related, albeit with lower support (83.6/58).

The PEBP protein family includes both promoters and repressors of flowering, which can be differentiated by key diagnostic amino acid residues (Ahn *et al*., 2006; Hanzawa *et al*., 2005; Ho and Weigel, 2014; Taoka *et al*., 2011; Wickland and Hanzawa, 2015). The protein sequence of CsFT1 was found to resemble the flowering promoters at key residues: CsFT1 possesses a tyrosine at position 134 and a tryptophan at position 138, while repressor FTs have non-tyrosine and non-tryptophan at these respective amino acid positions (Figure 5c).

### A PACE assay can be used for *CsFT1* genotyping in mapping populations and diverse hemp accessions

To facilitate selection for specific alleles through molecular breeding, PACE markers were developed targeting a SNP between the *CsFT1* ‘Felina 32’-type allele and both copies in the ‘FINOLA’-type versions of *CsFT1* (Table S3). The PACE assay was used on the previously sequenced ‘Felina 32’ and ‘FINOLA’ parent and F_1_ individuals of the mapping population, returning genotype calls expected on the basis of WGS data: homozygous for ‘Felina 32’-type allele, ‘FINOLA’-type allele and heterozygous, respectively. The PACE assay was further tested on plants cultivated under continuous light to test if allele call and flowering phenotype were associated. ‘FINOLA’ individuals flowered early and consistently were genotyped as heterozygous or homozygous ‘FINOLA’-type allele, while ‘Felina 32’ individuals grown in parallel, were flowering much later or not at all being predominantly genotyped as ‘Felina 32’- type or heterozygous (P = 5.244e^-5^, Mann–Whitney U-test) (Figure 6a). Subsequently, F_2_ individuals not previously sequenced were tested. Early flowering individuals (<60 d) were genotyped either heterozygous or homozygous for the ‘FINOLA’-type allele. Later flowering F_2_ individuals (>70 d) were mostly homozygous ‘Felina 32’-type, with some heterozygous, and two ‘FINOLA’-type allele individuals (Figure 6b). The allele call in the F_2_ is correlated to the flowering time phenotype, with ‘Felina 32’-type allele mean flowering time being significantly different from both heterozygotes and ‘FINOLA’-type allele (P = 4.9e^-6^, P = 3.2e^-8^, respectively, Kruskal Wallis with Conover-Iman post-hoc test). F_2_ plants genotyped as homozygous ‘FINOLA’-type or heterozygous were not significantly different from each other with respect to flowering time (P = 0.1932, Figure 6b). This suggests that the ‘FINOLA’-type allele may have a dominant effect on flowering phenotype.

**Figure 6.**
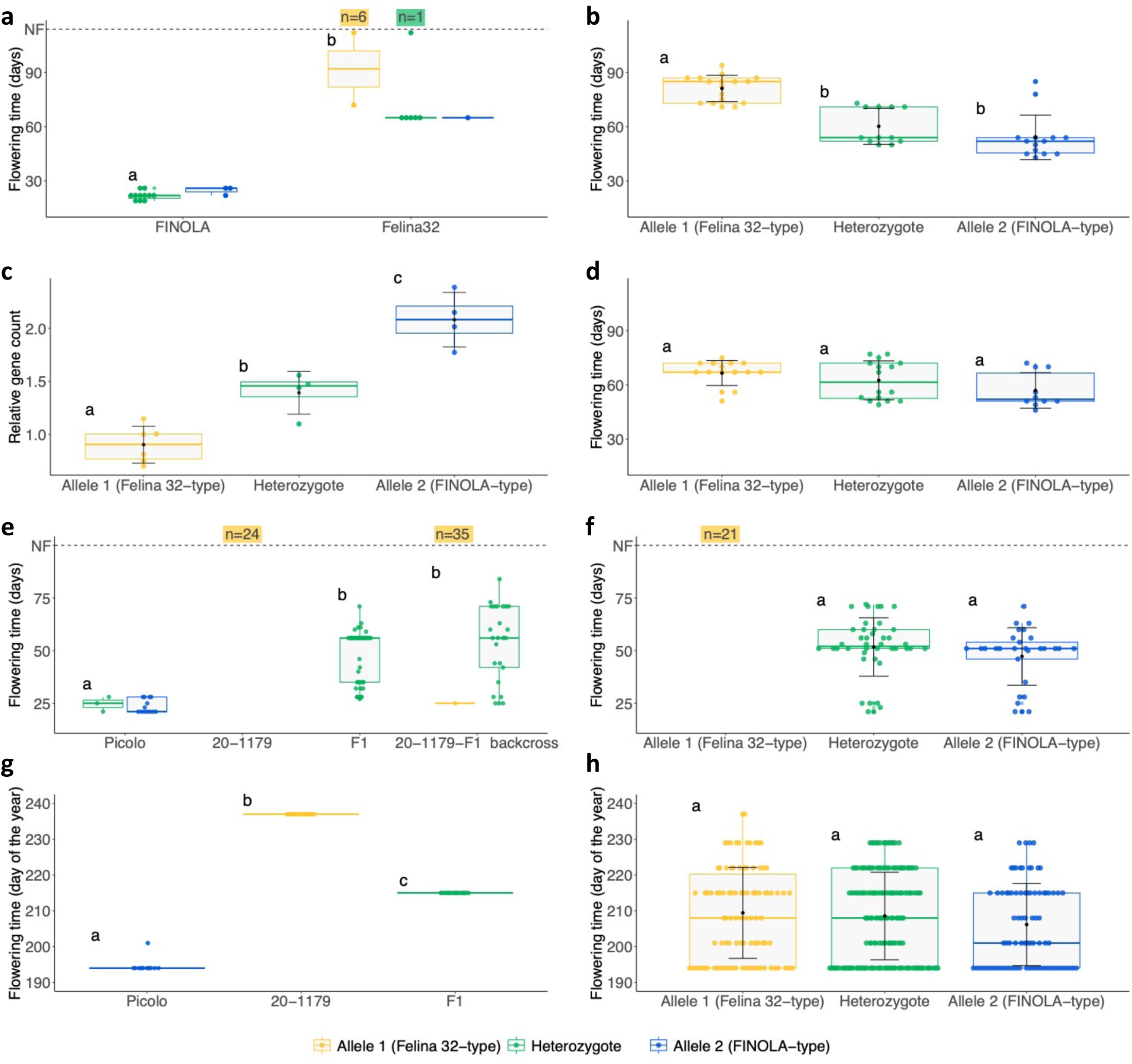
PACE-based genotyping of *CsFT1* in hemp accessions and mapping populations segregating for flowering time. A PACE assay was used to genotype individual *C. sativa* plants at the *CsFT1* locus. Individuals genotyped as ‘Felina 32’-type, ‘FINOLA’-type or heterozygous are depicted in yellow, blue or green, respectively. Flowering time and genotypes of female ‘FINOLA’ and ‘Felina 32’ individuals grown under continuous light. Flowering time of ‘FINOLA’ and ‘Felina 32’ was significantly different. Some ‘Felina 32’ individuals (n=7) did not flower (NF) by the conclusion of the experiment on day 112 and box colour denotes allele call of the NF plants (a). In the ‘Felina 32’ × ‘FINOLA’ F_2_ females grown in natural light conditions, allele call is significantly associated with the flowering time phenotype (b). *CsFT1* copy number and allele call are significantly associated, with ‘Felina 32’-type, heterozygotes and ‘FINOLA’-type alleles possessing a mean relative copy number of 0.9, 1.39 and 2.08 respectively (c). In the ‘Felina 32’ × ‘FINOLA’ F_3_ females grown in natural light conditions allele call is not significantly associated with flowering time (d). Flowering times under continuous light and genotypes for 20-1179, ‘Picolo’, the F_1_ and F_1_ backcross individuals (e). Flowering times were significantly different between ‘Picolo’ and both the F_1_ and F_1_ backcross. 100% of 20-1179 and 56.7% of the F_1_ backcross did not flower by the conclusion of the experiment on day 100. Flowering times of the F_2_ were significantly associated with allele call at *CsFT1* when grown under continuous light. No ‘Felina 32’-type allele plants flowered by the end of the experiment (100 days) (f). Flowering times of ‘Picolo’, 20-1179 and F_1_ individuals (g) and the F_2_ grown in the field (h). Here, allele call was not significantly associated with the flowering time. Non-flowering individuals were excluded from boxplots and statistical analyses. Mann–Whitney U-test (a, f) and Kruskal Wallis with Conover-Iman post-hoc test (b, c, d, e, g, h) were used. Statistical significance is reported at α = 0.05 and significance levels are indicated with letters.

We further tested whether allele call and the previously conducted copy number analysis based on qPCR can be correlated. The mean relative copy number of individuals with the ‘Felina 32’ allele was only half of the ones with the ‘FINOLA’ allele, which was a significant difference (0.9 vs. 2.08 respectively, P = 3.8e^-5^, Kruskal Wallis with Conover-Iman post-hoc test) (Figure 6c). Furthermore, the mean relative copy number of heterozygous individuals was in between, and significantly different from both homozygotes (mean = 1.39, P = 0.0052, P = 0.0074, respectively, Kruskal Wallis with Conover-Iman post-hoc test) (Figure 6c).

In the F_3_ population most early flowering individuals were genotyped as heterozygous or homozygous ‘FINOLA’-type and most later flowering individuals were genotyped as homozygous ‘Felina 32’-type allele (Figure 6d). However, groups with distinct homozygous allele types were not significantly different from one another with respect to flowering time (P = 0.0835, Kruskal Wallis with Conover-Iman post-hoc test) and the flowering time of heterozygotes was not significantly different from either homozygote (P = 0.4206, P = 0.2401, respectively, Kruskal Wallis with Conover-Iman post-hoc test) (Figure 6d).

Subsequently we tested the PACE assay on other cultivars and populations, including Cornell breeding pedigrees based on a cross between ‘Picolo’, an early flowering Canadian grain-type cultivar closely related to ‘FINOLA’ (Carlson *et al*., 2021), and 20-1179, a late flowering breeding line derived from a Chinese landrace (Figure 6e) to form a segregating population. Under continuous light, ‘Picolo’ and F_1_ plants as well as the majority of F_2_ plants flowered within 72 days (Figure 6e, f) with all of the early flowering plants genotyped homozygous or heterozygous for the ‘FINOLA’-type *CsFT1* allele. 20-1179 plants as well as some 23% of the F_2_ plants remained vegetative under continuous light until termination of the experiment 100 days after sowing. Those individuals all were homozygous for the ‘Felina 32’ allele of *CsFT1* (Figure 6e, f).

Interestingly, the ratio of flowering to non-flowering plants in this F_2_ population was very close to a 3:1 segregation (77% of the plants flowering, 23% non-flowering). This Mendelian segregation was further confirmed in a backcross of the F_1_ plants with the 20-1179 parent. Under continuous light 56.7% of the plants did not flower, which is close to the expected 1:1 segregation ratio for a Mendelian locus. All non-flowering plants were homozygous for the ‘Felina 32’ *CsFT1* allele (Figure 6e).

Under field conditions, ‘Picolo’ flowered very early (median flowering date July 13), 20-1179 flowered later (median flowering date August 25), and the F_1_ plants had an intermediate flowering time (median flowering date August 3) (Figure 6g). However, while there was segregation for flowering time in the F_2_ population (range = July 13-August 25), there was no association between flowering time and the *CsFT1* allelic status in the F_2_ under field conditions (Figure 6h, P = 0.22, Kruskal Wallis chi-squared test).

### The ‘FINOLA’-type allele of *CsFT1* is present in other early flowering hemp cultivars

To further assess the presence of the ‘FINOLA’-type *CsFT1* allele in various genetic backgrounds, the PACE marker was used to screen 50 *C. sativa* accessions (Carlson et al., 2021, Table S10). Accessions known to flower early in long day conditions such as ‘Picolo’, ‘CFX-2’, ‘Anka’, ‘Henola’ and ‘USO-31’ (Toth *et al*., 2022; Woods *et al*., 2021) were homozygous ‘FINOLA’-type or heterozygous (Figure 7a, Table S10). In contrast, accessions known to be photoperiod sensitive with ‘intermediate’ or ‘late’ flowering phenotypes such as ‘RN16’, ‘Late Sue’ and ‘PuMa’ (Stack *et al*., 2021; Toth *et al*., 2022), were all homozygous ‘Felina 32’-type allele (Figure 7a, Table S10). An accession named ‘Russian Auto CBD’ was homozygous for the ‘FINOLA’-type allele (Table S10). Several photoperiod-sensitive accessions, such as ‘A2R4’ and ‘Felina 32’, were homozygous or heterozygous for the ‘FINOLA’-type allele (Figure 7a, Table S10).

**Figure 7.**
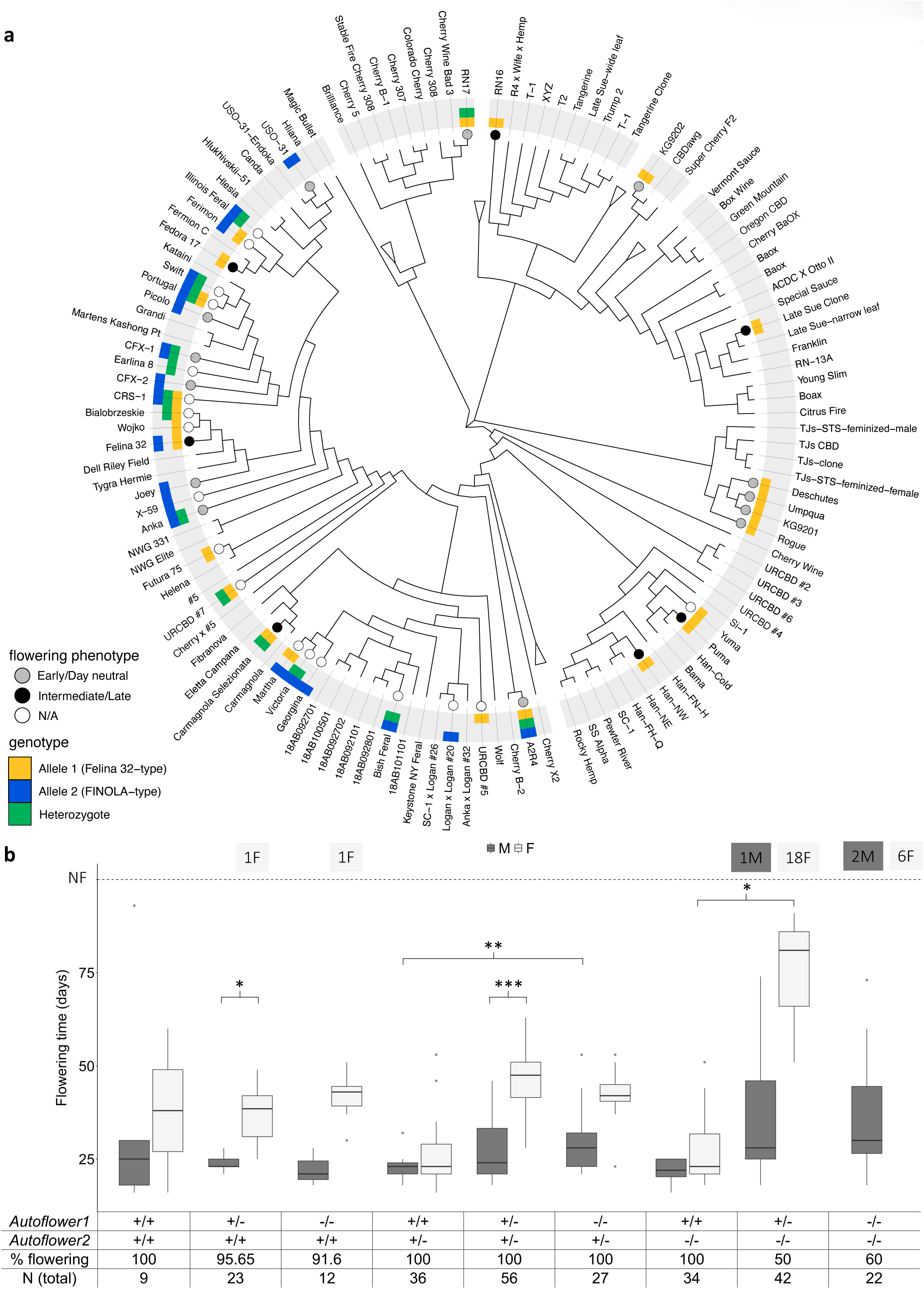
Photoperiod insensitivity has two independent origins in *C. sativa*. Genotyping results for a diverse panel of hemp accessions using the *CsFT1* PACE essay (a). Relationships between accessions are depicted by a maximum likelihood tree (Carlson *et al*., 2021). Blue, yellow and green squares in the outer circles depict a ‘FINOLA’-like, ‘Felina 32’-like or a heterozygous genotype, respectively. Grey, black and white filled circles at branch ends depict early/day-neutral, intermediate/late and unknown flowering time. Flowering time and genotypes of an F_2_ population formed by crossing ‘Le Crème’ (homozygous *Autoflower1, +/+ // -/-*) with ‘Picolo’ (homozygous *Autoflower2, -/- // +/+*) (b). Plants were grown under continuous light and genotyped using the PACE assay described here and previously (Toth *et al*., 2022). All statistical comparisons are based on the Kruskal Wallis with Conover-Iman post-hoc test and significance levels are indicated with asterisks (* p<0.05, ** p<0.0005, *** p<0.00001).

Some early flowering photoperiod-sensitive accessions such as ‘Umpqua’, ‘Deschutes’ and ‘Rogue’ (Stack *et al*., 2021; Toth *et al*., 2022) were homozygous for the ‘Felina 32’-type allele at *CsFT1*. Furthermore, some accessions that are known to be photoperiod-insensitive and homozygous *Autoflower1* (Toth *et al*., 2022), such as ‘AutoCBG’, KG9201 and KG9202 (Stack *et al*., 2021; Toth *et al*., 2022), were genotyped as homozygous ‘Felina 32’-type allele at *CsFT1* (Figure 7a, Table S10). Finally, as the PACE genotyping SNP is located within an exon, public RNA-seq datasets with high *CsFT1* expression were investigated, and we verified that the ‘FINOLA’ and ‘X-59’ samples possess the ‘FINOLA’ allele.

### A dihybrid cross between *Autoflower1* and *Autoflower2* shows a bias towards male flowering

Given the apparent distinction between *Autoflower1* and *Autoflower2* based on mapping location and marker calls we wanted to test the interaction between the two loci. For this purpose we created a population segregating for both loci by crossing a female individual of ‘Le Crème’ homozygous for *Autoflower1* (+/+ // -/-) with four ‘Picolo’ male individuals homozygous for *Autoflower2* (-/- // +/+).

In about 88% of the plants of the ‘Le Crème’ × ‘Picolo’ F_2_ population flowering was observed under continuous light. Flowering time was correlated with the genotype at *Autoflower1* and *Autoflower2* (Figure 7b).

On average, male individuals flowered earlier than female individuals of the same genotype, however, this was only significant for individuals heterozygous for *Autoflower1* and at least one ‘FINOLA’ allele at *Autoflower2* (Figure 7b). Furthermore the genotype at *Autoflower1* seems to affect flowering time of heterozygous *Autoflower2* plants.

Plants genotyped to be homozygous for the photoperiod-sensitive flowering allele in both *Autoflower1* and *Autoflower2* (-/- // -/-) had the greatest proportion of non-flowering individuals. In this group, flowering was still observed, however exclusively male individuals flowered.

A significant proportion (∼12%) of the F_2_ plants did not flower under continuous light (Figure 7b), less than the proportion expected for a Mendelian dihybrid cross with one recessive and one dominant trait (18.75%). Genotyping showed that most of the non-flowering plants were female, while only a small number were male (79% and 9%, Figure 7b).

## Discussion

### An *FT-like* gene is a candidate for *Autoflower2* photoperiod insensitivity in the hemp cultivar ‘FINOLA’

The acquisition of photoperiod insensitivity has been deployed for latitudinal adaptation for many different crops throughout domestication (Huang *et al*., 2018; Hung *et al*., 2012; Soyk *et al*., 2017; Weller *et al*., 2019). Here, we identified *Autoflower2* as a locus associated with photoperiod insensitivity in the hemp cultivar ‘FINOLA’ (Figure 1, 2). *Autoflower2* appears to be dominant given the ability of heterozygotes to flower under continuous light (Figure 6), but the segregation for flowering time within allelic groups suggests that there are additional genes involved in photoperiodic control of flowering. *Autoflower2* is distinct from the previously described *Autoflower1* on chr01 which was identified in a photoperiod-insensitive high cannabinoid-type hemp accession (Toth *et al*., 2022).

The *Autoflower2* locus is approximately 0.5 Mbp in size and the confidence interval includes several candidate genes that may be involved in photoperiodic control of flowering (Figure 4a). Among those candidate genes, *CsFT1,* an ortholog of the flowering time gene *FT* from *A. thaliana,* (Figure 4b, c, 5b) is of particular interest. *FT-like* genes are highly conserved flowering promoters (Wickland and Hanzawa, 2015). Members of this gene family have a recurring role in flowering adaptation in various species, including *Zea mays*, *O. sativa*, *Helianthus annuus*, *Solanum lycopersicum* and *Glycine max* (Blackman *et al*., 2010; Guo *et al*., 2018; Soyk *et al*., 2017; Wu *et al*., 2017; Jia Zhang *et al*., 2015).

Copy number variations in key flowering regulators are known to affect flowering time in *A. thaliana* and *Triticum aestivum* (Rosas *et al*., 2014; Würschum *et al*., 2018), and data from barley suggest that increased copy numbers of *FT*-like genes are associated with early flowering (Nitcher *et al*., 2013). Furthermore, in triploid hemp, decreased dosage of the dominant photoperiod-sensitive allele at *Autoflower1* results in earlier flowering (Kurtz *et al*., 2023). Therefore it is plausible that copy number variation of different genes in the photoperiod pathway can contribute to a photoperiod-insensitive phenotype. Our findings suggest that *CsFT1* is tandem duplicated in the photoperiod-insensitive cultivar ‘FINOLA’, giving rise to *CsFT1a* and *CsFT1b* (Figure 4b, c, S5). Hence, it is possible that increased gene dosage of *FT*-like genes in ‘FINOLA’ is an important factor for its photoperiod insensitivity.

Beyond the *CsFT1* gene duplication, we identified sequence differences between *CsFT1* and *CsFT1a* and *CsFT1b*. The coding sequences of the alleles contain multiple non-synonymous differences, which might confer functional variation between alleles as well as between the two genes in ‘FINOLA’ (Figure 5a). Furthermore, intronic sequences are highly dissimilar between *CsFT1*, *CsFT1a* and *CsFT1b* (Figure 4c). This could have an impact on the gene function due to, for example, the presence/absence of regulatory elements or splice sites (Kesari *et al*., 2012; Liu *et al*., 2021). Additionally, we identified a putative helitron-like transposable element in the second intron of *CsFT1a*, which is absent from both *CsFT1* and *CsFT1b* (Figure S7). It is well established that transposon insertions can impact transcription and splicing patterns (Gill *et al*., 2021). Transposon insertions in introns of *FT*-like genes in *G. max* and *Brassica rapa* caused reduced transcription and consequently later flowering (Wu *et al*., 2017; Xueming Zhang *et al*., 2015; Zhao *et al*., 2016). In *A. thaliana*, extensive natural variation at the *FLOWERING LOCUS C* locus can be attributed to transposon insertions facilitating local adaptation regarding vernalization (Quadrana, 2020). Transposon insertions impacting regulatory and coding regions of three flowering time loci contributed to the northern expansion of *Z. mays* (Castelletti *et al*., 2014; Huang *et al*., 2018; Yang *et al*., 2013). In *Z. mays*, quantitative disease resistance was conferred by alternative splicing of a helitron-derived exon replacing that of the wild-type (Liu *et al*., 2020).

In the analysed RNA-seq, *CsFT1* is expressed in flowers, leaves, bast fibres, stem, roots and seeds (Figure 3). This expression pattern broadly follows homologs in other species such as *A. thaliana* (AT1G65480, Fernández et al., 2016)), *Z. mays* (GRMZM2G179264, Guo *et al*., 2018) and *G. max* (Glyma.02G069500, Wu *et al*., 2017) (Klepikova *et al*., 2016; Libault *et al*., 2010; Sekhon *et al*., 2011; Sullivan *et al*., 2019). Interestingly, among the tissues with the highest *CsFT1* expression are feral North American *C. sativa* flowers, ‘FINOLA’ flowers and seeds from the accession ‘X-59’ (Figure 3, Table S1).

Taken together, we propose that structural differences in an *FT*-like gene at the *Autoflower2* locus could be a major cause for the photoperiod insensitivity observed in ‘FINOLA’. The cause for the large structural variation and sequence differences between ‘FINOLA’ and ‘Felina 32’ is not entirely clear, but it is interesting to note that the genes are located at the end of the chromosome, which is a region that is gene rich and experiences high recombination frequencies (Laverty *et al*., 2019). In other plants like *T. aestivum* or *Phaseolus vulgaris*, recombination is also concentrated towards the chromosome ends and has been postulated to facilitate rapid adaptation to new environmental conditions (Chen *et al*., 2018; Glover *et al*., 2015; Schilling *et al*., 2020; IWGSC *et al*., 2018). It is possible that a similar mechanism is at play in *C. sativa* and that rapid evolution of an *FT* ortholog, due to its distal location on the chromosome, contributed to adaptation of hemp to different latitudes.

### Photoperiod insensitivity originated multiple times independently in *C. sativa*

Several flowering time QTL have been identified in *C. sativa*, with causal genes and mechanisms remaining undetermined (Petit *et al*., 2020; Toth *et al*., 2022; Woods *et al*., 2021). A GWAS for flowering time identified 28 candidate genes related to flowering, including three *PEBP*-like genes *CsCEN1* (LOC115697843), *CsFT2* (LOC115709185) and *CsFT3* (LOC115697736) (Petit *et al*., 2020). We confirmed the presence of three other *FT*-like genes, *CsFT2*, *CsFT3* and *CsFT4* in the ‘CBDRx’ reference genome (Figure 5b).

*Autoflower1* and *Autoflower2* are clearly functionally and phylogenetically distinct from each other. *Autoflower1* and *Autoflower2* are located on chr01 and chr08, respectively. They fail to complement each other in segregating F_2_ populations which include non-flowering plants when grown under continuous light (Figure 7b). While *Autoflower1* is recessive (Toth *et al*., 2022), *Autoflower2* is dominant with respect to flowering under continuous light. Markers developed for breeding were tested on a diverse germplasm and further confirm the distinction between *Autoflower1* (KG9202, Toth et al., 2022) and *Autoflower2* (‘FINOLA’ and ‘Picolo’) in conferring photoperiod insensitivity (Figure 7a). This suggests that photoperiod insensitivity originated multiple times independently in *C. sativa*. Interestingly, ‘Russian Auto CBD’ appears to be photoperiod-insensitive through *Autoflower2*, not *Autoflower1* (Table S10). ‘Russian Auto CBD’ is also in the same clade as ‘FINOLA’ as determined through GBS (Carlson *et al*., 2021, unpublished data). Like ‘FINOLA’, ‘Russian Auto CBD’ is reported to have been selected from Russian landraces (Callaway and Laakkonen, 1996; https://www.nativcanna.at/shop/hanfsamen/russian-auto-cbd/), which may indicate that *Autoflower2* was derived from the putative species of *Cannabis ruderalis*, which is theorised to have originated in Siberia (Callaway and Laakkonen, 1996; Toth *et al*., 2022). Further work is required to investigate the taxonomic origins of both *Autoflower1* and *Autoflower2*, but the independent origins of photoperiod insensitivity supports the likelihood of a complex domestication history in *C. sativa*.

Two recent mapping studies have suggested *C. sativa* homologs of *APETALA2* and *PSEUDO-RESPONSE REGULATOR 37* (*PRR37*) as candidate genes within *Autoflower1* (Garfinkel *et al*., 2023; Leckie *et al*., 2023). Alterations of *PRR37* homologs in *T. aestivum*, *O. sativa* and *Sorghum bicolor* are known to confer photoperiod insensitivity due to de-repressed *FT* expression (Beales *et al*., 2007; Koo *et al*., 2013; Murphy *et al*., 2011). It is thus conceivable that different genes of the photoperiod pathway were affected independently of each other to bring about photoperiod insensitivity in *C. sativa*. Similar observations have been made in *G. max*, where mutations in different genes of the photoperiod pathway can lead to reduced photoperiod sensitivity (Cao *et al*., 2017; Xu *et al*., 2013).

In dioecious *C. sativa,* males typically flower before females (protandry) (Figure S2b) (Nelson, 1944; Salentijn *et al*., 2019; Toth *et al*., 2022). The genetic control of protandry in *C. sativa* is unknown. In all dioecious accessions and all populations created in this study, male individuals consistently flowered earlier than female individuals and monoecious individuals (Figure S2, 7b). Interestingly, the vast majority of non-flowering individuals in the ‘La Creme’ × ‘Picolo’ F_2_ population were females (Figure 7b). *Vice versa*, flowering could be observed in plants that possessed the photoperiod sensitive alleles at *Autoflower1* and *Autoflower2* (i.e. a -/- // -/- genotype). However, here only male individuals were flowering. This is an indicator of sexually dimorphic photoperiod-insensitive flowering in *C. sativa*, which may involve an interaction with a locus on the Y chromosome.

### Genetic and environmental factors may determine the significance of *Autoflower2* in controlling flowering time

While *Autoflower2* appears to convey photoperiod insensitivity, its effect on flowering time is not always consistent. In the ‘Felina 32’ × ‘FINOLA’ F_3_ generation analysed here, the *Autoflower2* PACE markers were not significantly associated with flowering time (Figure 6d). This is in contrast to the F_2_, where a strong association was detected. The glasshouse temperatures in 2021, the season when the F_3_ was grown, were considerably higher than for 2020, the F_2_ growth season (Figure S1). It is known that temperature is an important factor in controlling flowering time in *C. sativa* (Cosentino *et al*., 2012; Hall *et al*., 2012; Lisson *et al*., 2000; Nelson, 1944). This notion is supported by the observation that later flowering individuals in the F_3_ population were flowering approximately 11 days earlier than the late flowering individuals of the F_2_ population. In other words, flowering of the F_3_ population seemed to be accelerated, possibly due to higher temperatures (Figure S1). In *A. thaliana* flowering can be induced by high temperatures under non-inductive short-day conditions due to activation of *FT* (Balasubramanian *et al*., 2006; Fernández *et al*., 2016; González-Suárez *et al*., 2023). In *G. max* the photoperiod pathway can be circumvented under inductive short-day conditions as temperatures up to 30°C accelerate flowering via *FT* expression, whereas temperatures above 35°C can interact with the photoperiod to delay flowering (Tang *et al*., 2022). It is possible that a similar, complex regulatory pathway exists in *C. sativa*, and that the acceleration of flowering time at higher temperatures masks the effect of photoperiodic control to some extent. This would explain why association between *Autoflower2* and flowering time was not significant in the F_3_ generation. In the 20-1179 × ‘Picolo’ F_2_, *Autoflower2* allelic calls fully explained flowering patterns under continuous light, and flowering time seemed to follow a Mendelian segregation. In contrast, *Autoflower2* allelic calls were not associated with flowering time under field conditions (Figure 6g, h). This further suggests a role for environmental factors in controlling flowering time in addition to *Autoflower2*, possibly due to factors intrinsic to glasshouse culture including limited root space, a known factor in affecting flowering time (Schilling *et al*., 2023).

Varying day lengths may also determine the significance of *Autoflower2* in controlling flowering time: under continuous light *Autoflower2* behaved like a Mendelian locus controlling flowering time (Figure 6e, f). In contrast, in the ‘Felina 32’ × ‘FINOLA’ F_2_ which was grown under long summer days (latitude 53.305° N) *Autoflower2* did have a quantitative, but not Mendelian effect on flowering time (Figure 1, 6b). Finally, in the 20-1179 × ‘Picolo’ F_2_ grown in the field under shorter summer days (latitude 42.869422 N) no significant effect of *Autoflower2* on flowering time was observed (Figure 6h).

It is also noteworthy that an individually sequenced female ‘FINOLA’ plant did not carry a *CsFT1* duplication (Figure 4c), suggesting the cultivar is not genetically uniform at the *Autoflower2* locus. This is in line with the variability observed in the results of the qPCR-based copy number variation analysis for ‘FINOLA’ and F_2_ individuals, which displayed gene counts in line with 1 copy, 2 copies or a heterozygous state equivalent to 1.5 genes (Figure 4d). Many *C. sativa* cultivars are not genetically uniform, and cultivars of the same name but from different sources can behave differently under the same environmental conditions (Zhang *et al*., 2021).

Based on PACE genotyping, both ‘FINOLA’ and ‘Felina 32’-like alleles of the *Autoflower2* locus seem to exist in other accessions (Figure 6, 7a). In some of the accessions categorised as ‘Intermediate/Late’ flowering such as ‘A2R4’ and ‘Felina 32’ both alleles of *CsFT1* could be detected (Figure 7). In these accessions some individuals will eventually flower under long-day conditions (Stack *et al*., 2021; Toth *et al*., 2022) and so segregation at this locus may explain this phenotypic variability. It will be interesting to further test how exactly flowering times vary within those cultivars, whether the allelic status of *Autoflower2* is correlated with variation in flowering time, and to what extent environmental variation other than photoperiod affect flowering time.

Together, our data indicate that there is a complex interplay between different flowering time pathways in *C. sativa*, and whether or not particular *Autoflower2* alleles accelerate flowering may depend on various environmental conditions.

### *ELF9* and *AGO5*-like genes co-locate with *Autoflower2* and are candidate genes for photoperiod insensitivity

While the structural variant in *CsFT1* is a strong causal candidate for the photoperiod-insensitive phenotype, the genes *CsELF9*, *CsAGO5a* and *CsAGO5b* are in close vicinity to *CsFT1* and also are located within the *Autoflower2* locus.

In *A. thaliana*, ELF9 is an RNA-binding protein that targets transcripts of the central flowering regulator *SUPPRESSOR OF OVEREXPRESSION OF CONSTANS 1* (*SOC1*). *ELF9* and its homologs have been identified as candidate genes for flowering time in *A. thaliana* and *Z. mays* (Hung *et al*., 2012; Song *et al*., 2009). ELF9 from *A. thaliana* is a repressor of flowering and may reduce *SOC1* mRNA levels via nonsense-mediated decay (Song *et al*., 2009). The *elf9* mutant *A. thaliana* plants possessed a strongly reduced photoperiod sensitivity, similar to the phenotype of ‘FINOLA’.

In the analysed RNA-seq, *CsELF9* was highly expressed in all tissues (Figure 3). For the *CsELF9* homologs previously implicated as flowering time candidate genes in *A. thaliana* (AT5G16260, Song *et al*., 2009) and *Z. mays* (GRMZM2G353114, GRMZM2G074351, GRMZM2G171660, Hung *et al*., 2012), a similar expression pattern was observed whereby expression was observed throughout all tissues (Klepikova *et al*., 2016; Sekhon *et al*., 2011; Sullivan *et al*., 2019).

The *CsELF9* alleles of ‘FINOLA’ and ‘Felina 32’ possess a number of differences in CDS, and though all non-synonymous changes are outside of the regions encoding for the RNA recognition motif (Figure S4a) we cannot exclude the possibility that the differences are of functional relevance. However, in *A. thaliana*, *elf9* is a recessive loss of function mutation, whereas *Autoflower2* is dominant (Figure 6b). Thus, although we cannot exclude that *CsELF9* is the causal gene for photoperiod insensitivity in ‘FINOLA’, the dominance observed in the F_1_ population makes *CsFT1* a more likely candidate.

*AGO5* mediates miRNA156 activity in the meristematic tissue of *A. thaliana* (Roussin-Léveillée *et al*., 2020). Low miRNA156 levels in *ago5* results in early flowering (Roussin-Léveillée *et al*., 2020). *AGO5* in *A. thaliana* may be associated with transposon expression (Bradamante *et al*., 2022). AGO5 proteins have three known functional domains, PAZ, MID and PIWI (Mallory and Vaucheret, 2010). Between ‘FINOLA’ and ‘Felina 32’ we observed several amino acid changes in the predicted protein sequences of CsAGO5a and CsAGO5b that occur in these domains, potentially impacting protein function (Figure S5). Similar to *ELF9*, *AGO5* is a repressor of flowering, and at least in *G. max* loss of function mutations of the *AGO5* homolog are recessive (Cho *et al*., 2017), which is contrary to what we observe for *Autoflower2*.

Both *CsAGO5a* and *CsAGO5b* were weakly expressed in various tissues (Figure 3). Similarly, *AGO5* homologs involved in gamete development and flowering in *A. thaliana* (At2g27880, Roussin-Léveillée *et al*., 2020; Tucker *et al*., 2012) and seed colour in *G. max* (Glyma.11G190900, Cho *et al*., 2017) were expressed at a low level across most tissues (Klepikova *et al*., 2016; Libault *et al*., 2010; Sullivan *et al*., 2019). An exception to this is that in *A. thaliana* AGO5 can be highly expressed in flowers, a pattern absent in the analysed *C. sativa* data (Figure 3). In *A. thaliana*, *AGO5* is not involved in photoperiod-dependent flowering but in the age-dependent flowering pathway (Roussin-Léveillée *et al*., 2020). However, functions of *AGO5*-like genes in other species are very diverse (Bradamante *et al*., 2022; Cho *et al*., 2017; Tucker *et al*., 2012), and it is possible that the ‘FINOLA’ allele of *CsAGO5* confers photoperiod sensitivity.

### Is *Autoflower2* a flowering time “supergene”?

*CsFT1*, *CsELF9* and *CsAGO5* could all be related to key phenotypes of the ‘FINOLA’ cultivar: *CsELF9* could be associated with earlier maturity; *CsAGO5* could have a role in age-related flowering; *CsFT1* could be the central integrator of flowering time with variants that result in photoperiod-insensitive flowering. Considering the close proximity and linked nature of the four candidate genes at *Autoflower2*, there is also the possibility that all of them together contribute to photoperiod insensitivity.

“Supergenes” have been reported for complex phenotypes like flowering time and heterostyly in *Mimulus* and *Primula,* as well as social behaviour in fire ants and mimicry in butterflies (Huu *et al*., 2020; Schwander *et al*., 2014). Moreover, it was recently reported that core genes of the circadian clock are in tight linkage with each other over large evolutionary distances, and that this linkage ensures the correct inheritance of circadian states across generations (Michael, 2022). In a similar manner, the close linkage of *CsELF9*, *CsAGO5* and *CsFT1* may ensure the inheritance of a photoperiod-insensitive phenotype. It will be interesting to further disentangle the *Autoflower2* locus and to analyse the contribution of gene regulatory circuits to photoperiod insensitivity in more detail.

## Data availability

All raw sequencing files generated in this study are available on NCBI Sequence Read Archive (SRA) under BioProject accession number PRJNA956491.

## Acknowledgements

CAD is supported by an Irish Research Council–Environmental Protection Agency Government of Ireland Postgraduate Scholarship (grant no. GOIPG/2019/1987) and Fulbright Ireland-Environmental Protection Agency Scholarship. JS is supported by a Chinese Research Council Postgraduate Scholarship (grant no. CSC 201908300031). This work was partially supported by a grant from the Foundation for Food and Agriculture Research Hemp Research Consortium with matching funds provided by Scotts Corporation and by a sponsored research agreement with Thai Leaf. We are grateful to McKenzie Schessl, Alexander Wares, and Elisabeth Porschet for their invaluable technical assistance, to Verve Seed Solutions for kindly providing seeds of ‘Picolo’ and to Joseph Calderone of Emplant DNA for the generous donation of hemp seed. We are grateful to members of the Flower Power lab and colleagues at the School of Biology and Environmental Science at University College Dublin (UCD) for useful discussions and technical support, in particular Bredagh Moran, Conor Whelan, Grace Pender, Rachita Pradhan, Seán O’Leary, Ifeoma Agu and Matthew Priestley.

## Notes

### Competing Interest Statement

The authors have declared no competing interest.

