## Supplementary Materials for "A *FLOWERING LOCUS T* ortholog is associated with photoperiod-insensitive flowering in hemp (*Cannabis sativa* L.)"

### Supplementary methods- QTL-seq pipeline

The QTL-seq pipeline was conducted using ‘Felina 32’ as Parent 1, and the ‘CBDRx’ genome sequence as reference. SNPs required a minimum depth coverage of 9 to be considered in downstream analyses. Sliding window analysis (1-Mb interval with 100-kb increment) occurred based on the average SNP index and Δ(SNP-index). Modified scripts are available on github (https://github.com/Caroline-Dowling/F1_QTL-seq_Itoh-et-al-2019_sonic_2021).

Considerable differences exist between the ‘CBDRx’ and Laverty ‘FINOLA’ genome assemblies (Prentout *et al.*, 2020). A faster QTL-seq pipeline was employed to repeat QTL mapping but using the publicly available ‘FINOLA’ genome (Laverty *et al.*, 2019) as a reference (Sugihara *et al.*, 2022) (Figure S3). To determine the unassembled Laverty ‘FINOLA’ contigs that coincide with the end of the ‘CBDRx’ chr08, the following was conducted; in the Galaxy platform, megablast was employed to blast both Laverty ‘FINOLA’ and ‘CBDRx’ genomes with the ‘CBDRx’ coding sequences (https://ftp.ncbi.nlm.nih.gov/genomes/all/GCF/900/626/175/GCF_900626175.2_CBDRx/GCF_900626175.2_CBDRx_cds_from_genomic.fna.gz, last accessed on 23 Sept. 2019). The blast outputs were then intersected to create a table of the blast hits side by side, which was used as input in shinyCircos (Yu *et al.*, 2018) (Figure S3). The QTL-seq analysis was repeated with the 9 unassembled ‘FINOLA’ contigs above 100 kb in length that correspond to the end of chr08 in ‘CBDRx’, with a sliding window of 100 kb in increments of 10 kb (Figure S3).

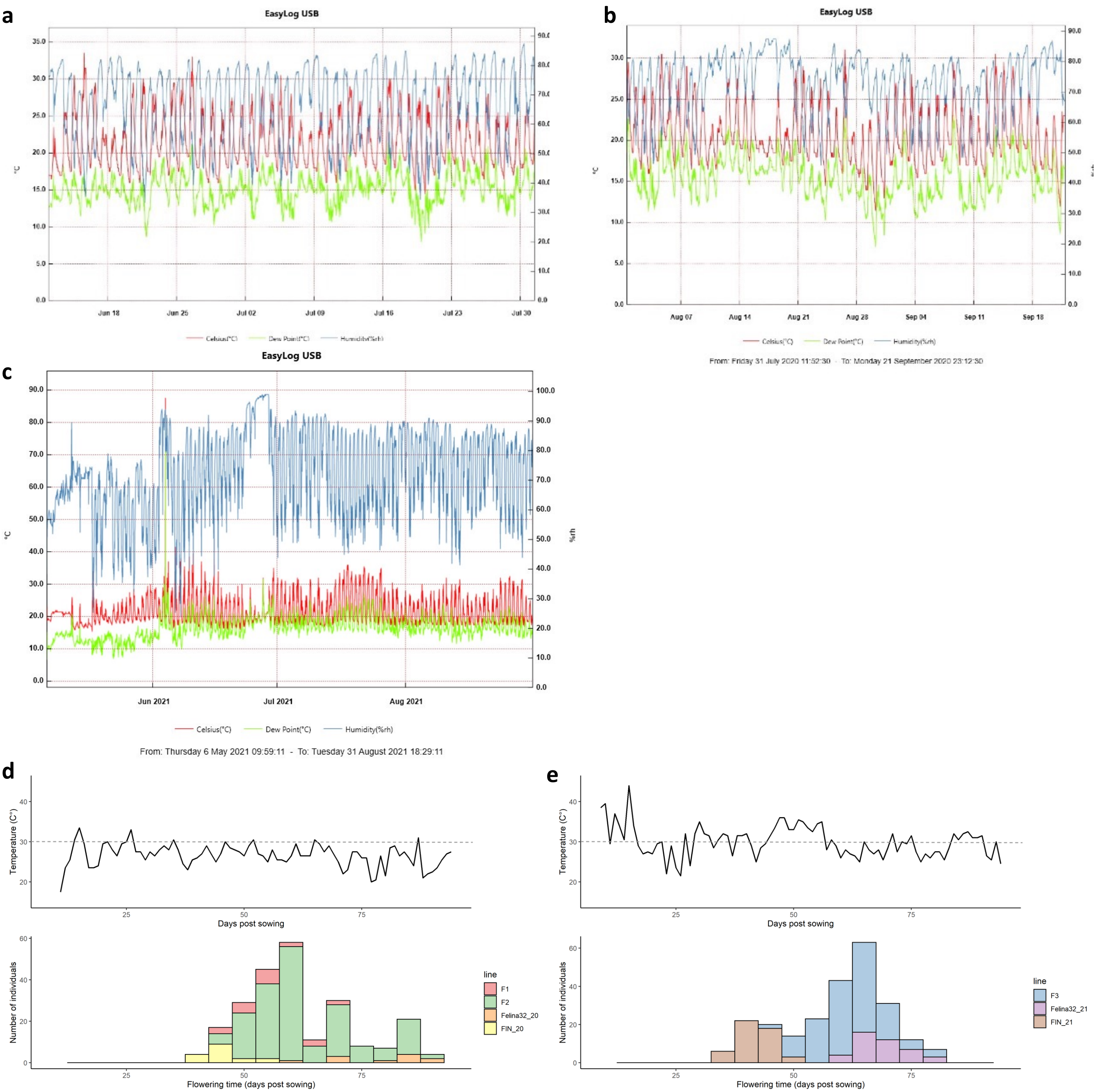


**Figure S1.** Temperature, dew point, humidity data records in 2020 (a) (b) and 2021 (c). The highest daily temperature in the glasshouse during plant cultivation, and the flowering time of female plants in (d) 2020 and (e) 2021. Higher temperatures were observed in 2021 and may have influenced flowering times.

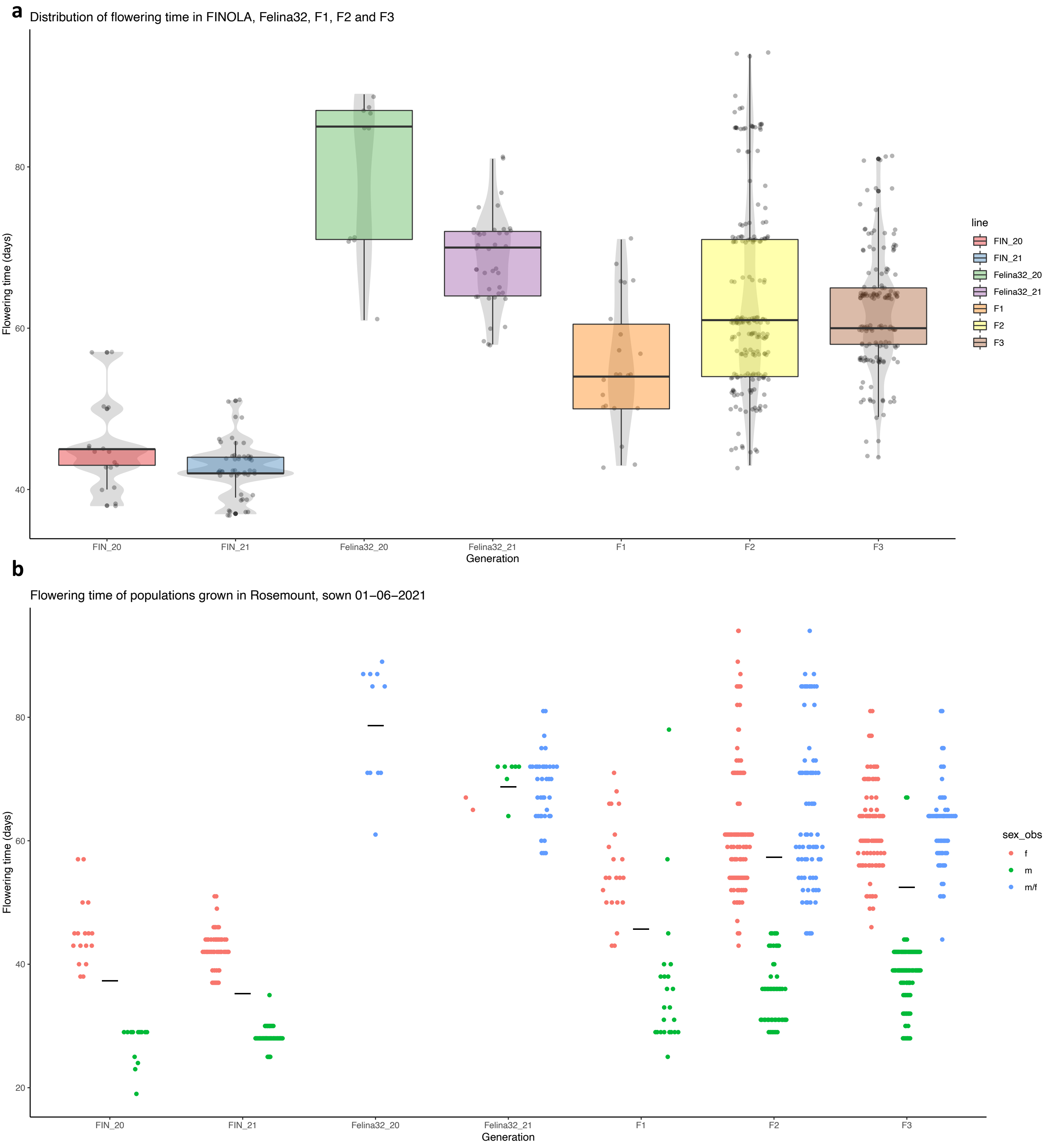


**Figure S2. Flowering time phenotyping data across several years.** (a) Flowering of ‘FINOLA’, ‘Felina 32’ in 2020 and 2021, $F_1$, $F_2$ and $F_3$ populations. (b) Sex related differences in flowering time of individuals grown under natural long day conditions in the glasshouse in Ireland. Males flower earlier than females and monoecious individuals in all populations.

**a**

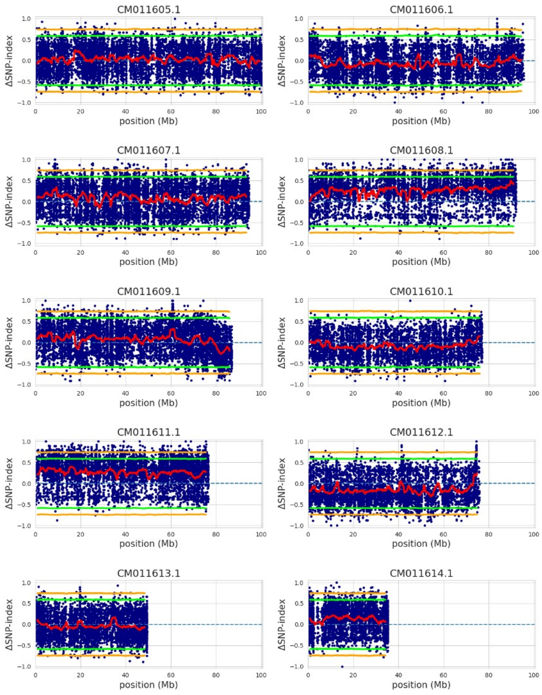


**c**

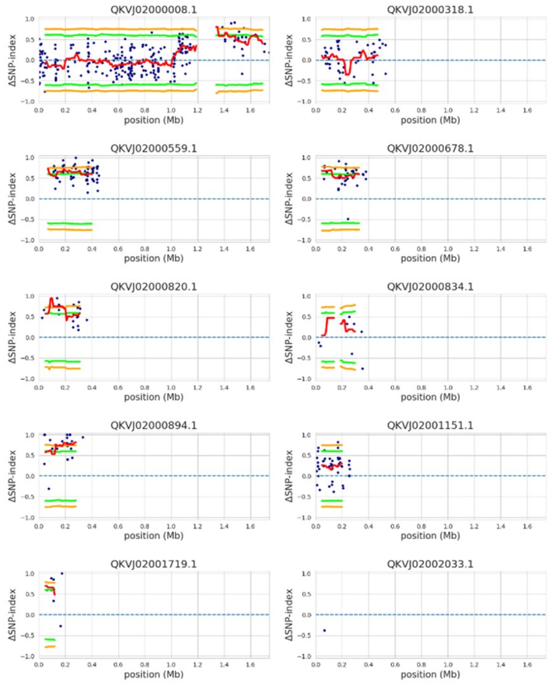


**b**

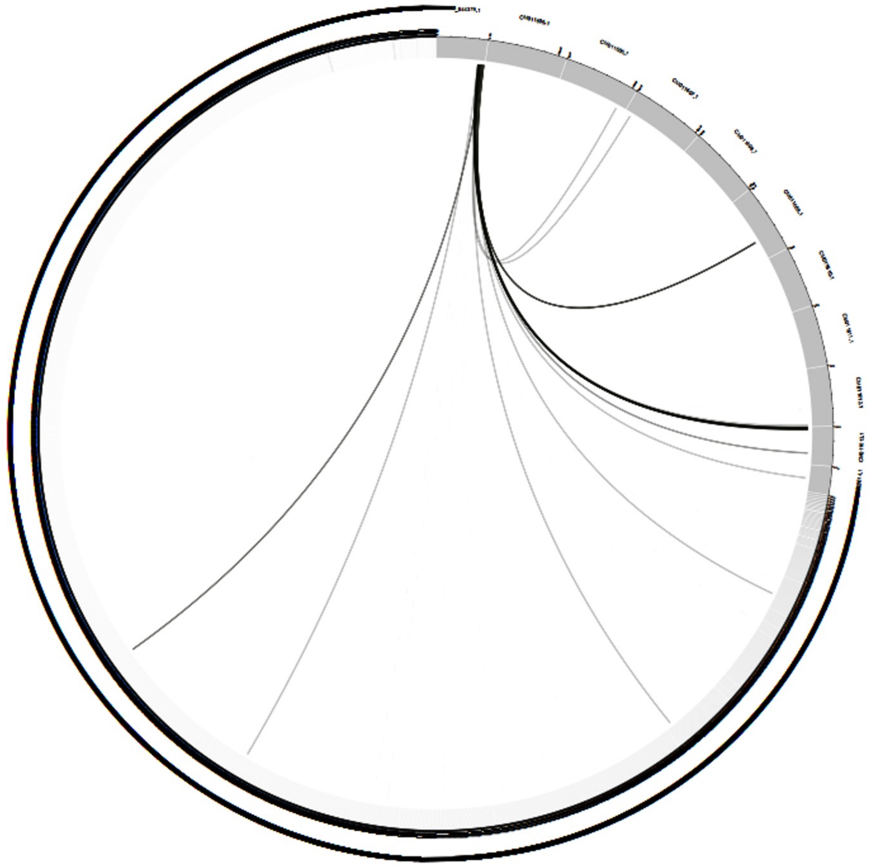

**Figure S3. QTL-seq analysis when 'FINOLA' (Laverty et al., 2019) is used as a reference genome.** (a) QTL-seq using 'FINOLA' Laverty genome as reference. No significant hit was detected on the chromosomes. (b) Circos plot comparing the end of chr8 in the 'CBDRx' genome to the Laverty assembly. This method identified the unassembled contig in the Laverty FINOLA genome where CsFT1 resides. (c) Repeated QTL-seq (using 'FINOLA' Laverty genome as reference) with contigs above 100kb in length. 6 out of the 9 contigs have a significant hit. Some hits are likely due to small size in the sliding window analysis, and so only contig QKVJ02000894.1 was analysed in further detail.

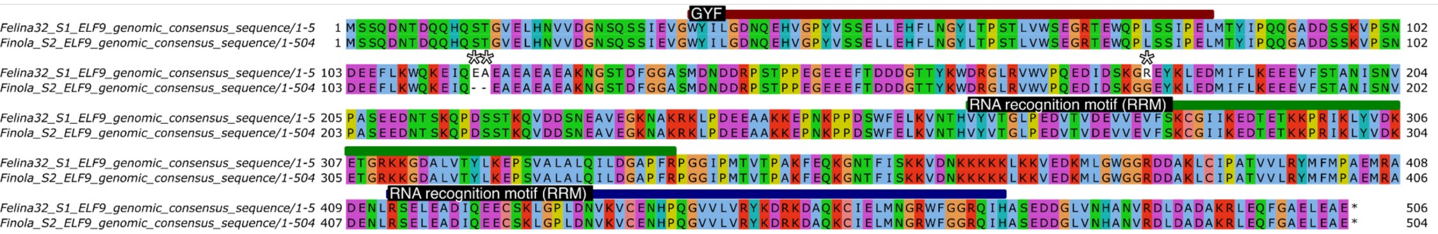


**Figure S4. Protein sequence alignment of candidate gene *CsELF9* in the chr08 QTL.** In CsELF9 (LOC115699158), a heterozygous (21 reads with deletion, 5 reads without) two amino acid deletion in a repeat in 'FINOLA' and one amino acid substitution between 'FINOLA' and 'Felina 32' are present outside of the known protein domains. To obtain coding sequences, genomic sequences were aligned to 'CBDRx' mRNA annotation in MAFFT, and UTRs and introns were removed in Jalview (Katoh et al., 2019; Waterhouse et al., 2009). If ambiguous nucleotides were present, the sequence was altered to the base with greater read support. If indels were present in coding sequences, sequences were altered as the IGV consensus sequence tool does not include indels. An indel was deemed genuine if present in genomic mappings to both reference genomes ('CBDRx', GCF_900626175.2 and FINOLA, GCA_003417725.2). Coloured bars signify protein domains and white stars denote positions with amino acid changes and dots highlight sequence gaps.

a

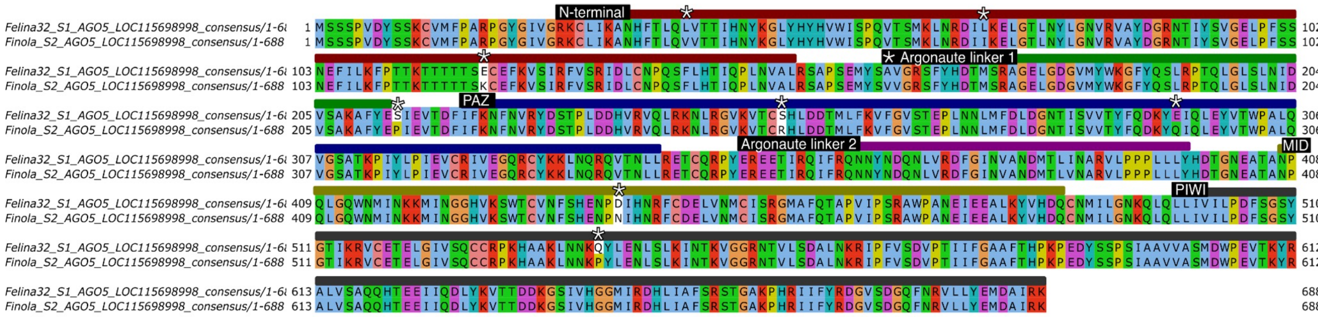


b

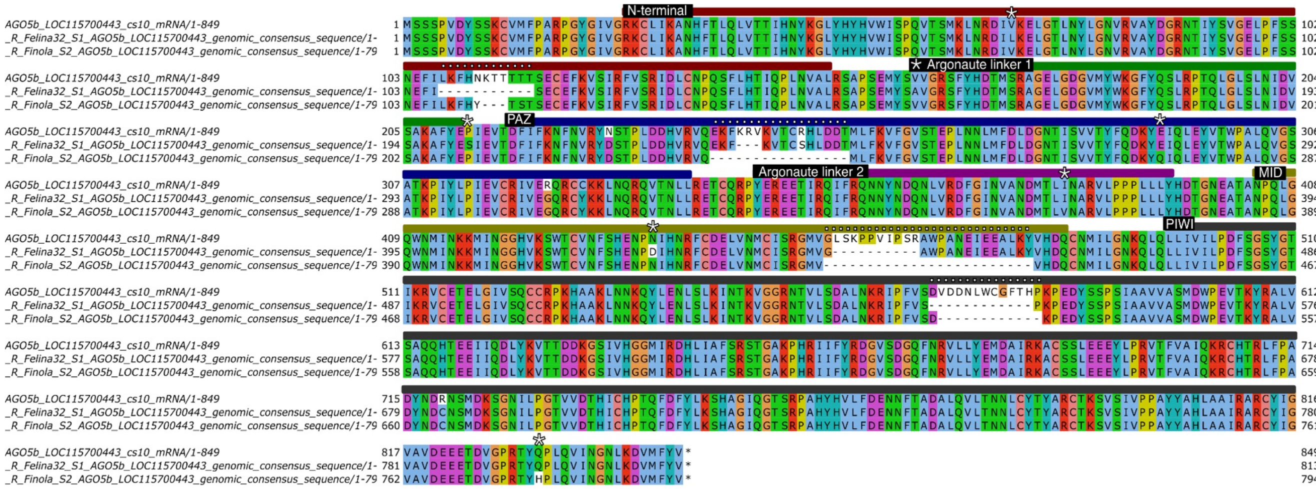


**Figure S5. Protein sequence alignments of candidate genes AGO5a and b in the chr08 QTL. Figure S5.** CsAGO5a (LOC115698998) differs in nine amino acid positions between 'FINOLA' and 'Felina 32', of which eight are in protein domains (a). CsAGO5b (LOC115700443), seven amino acids vary, with six differences located in protein domains (b). We note that *CsAGOa* and *b* sequences are very similar and might be misassembled as two separate genes. When comparing 'FINOLA' and 'Felina 32', five amino acid changes are shared between AGO5a and b. In CsAGO5b, four gaps (-) in the protein sequence are present in both 'FINOLA' and 'Felina 32' when compared to CBDRx, caused by ambiguous nucleotides ('Ns') due to a lack of reads mapping to that region. Both *CsAGOa* and *b* are located beside unannotated regions (schematic of locus, Figure 3b). Supplemental nucleotides have been added to the end of *CsAGO5a* in the reference genome as it is at the chromosome end, hence the last 163AA of the protein were not analysed as both 'FINOLA' and 'Felina 32' sequencing data aligned poorly to this region (see NCBI https://www.ncbi.nlm.nih.gov/gene/?term=LOC115698998). For the alignment method see Figure S4 legend.

**a**

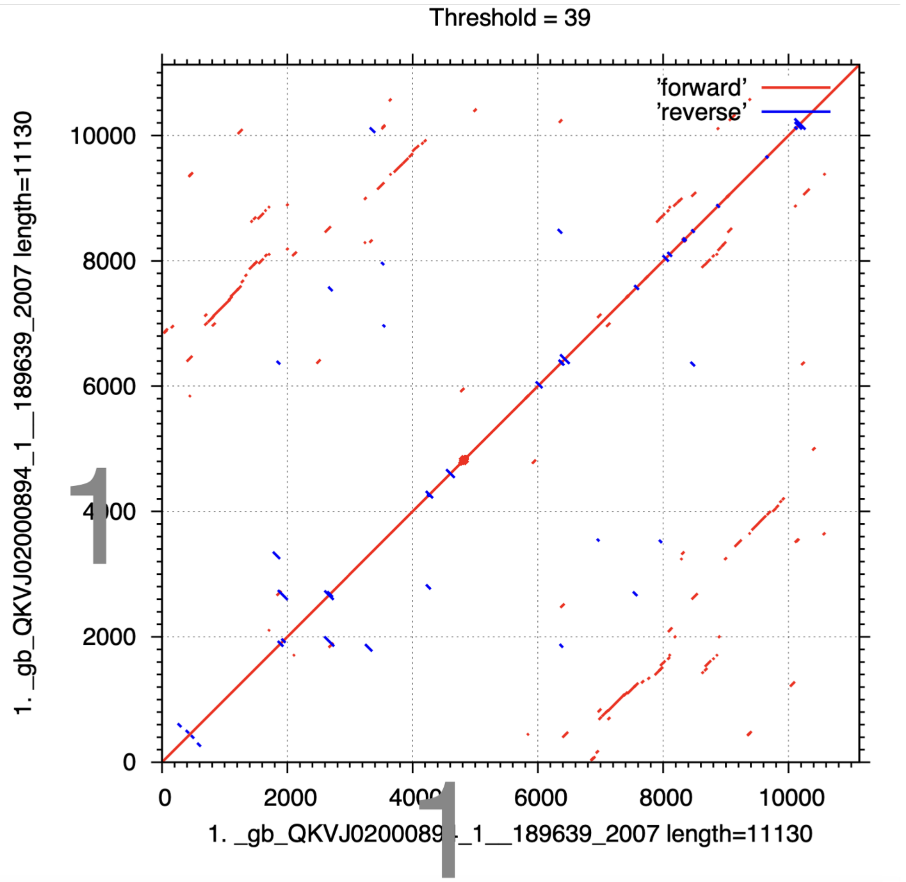


**b**

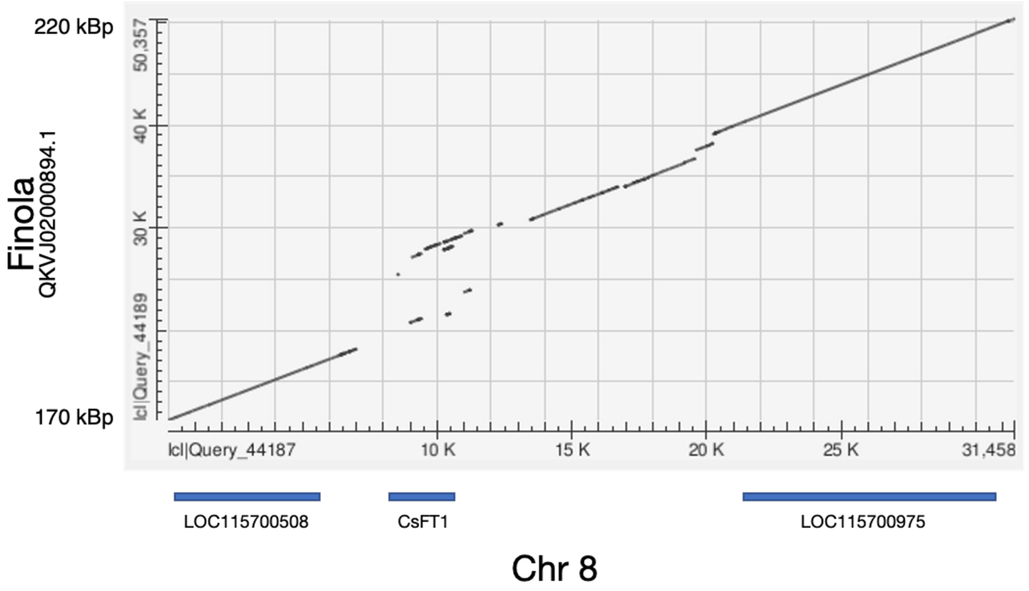


**Figure S6. *CsFT1* locus in FINOLA and CBDRx.** (a) Dot plot of the 11 kb region (QKVJ02000894.1:189639-200768) on the contig QKVJ02000894.1 on which the *CsFT1* duplication is found in the 'FINOLA' genome (GCA_003417725.2). The exons of *CsFT1* copies are clearly in duplicate on the contig. (b) Dot plot of the 'FINOLA' contig QKVJ02000894.1 compared to the chr08 region in which *CsFT1* is located in the 'CBDRx' genome assembly. The genes upstream and downstream of *CsFT1* are in synteny with the 'FINOLA' contig, suggesting only the *CsFT1* genes are impacted by this structural variant.

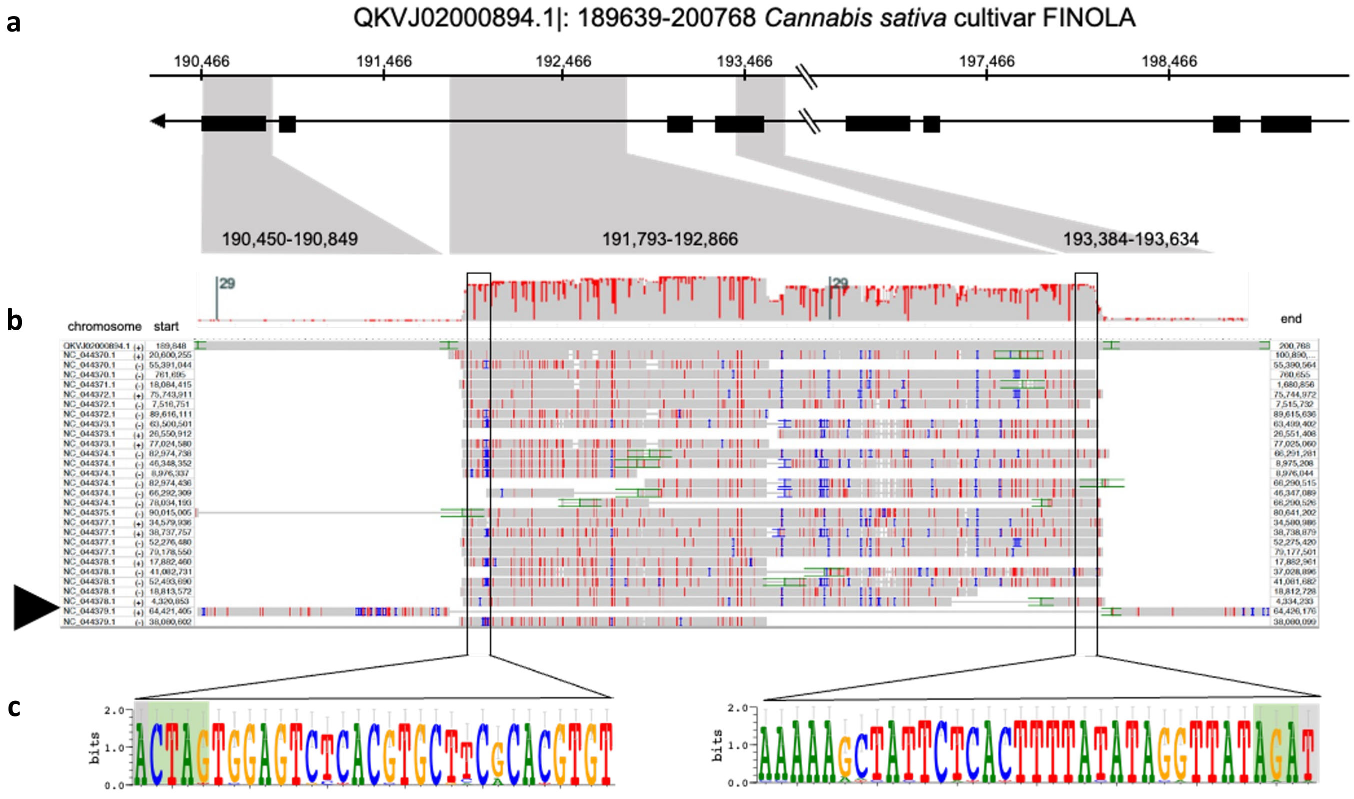


**Figure S7. A repetitive element with helitron-like sequence motifs is located in the large intron of *CsFT1a* in 'FINOLA'.** The 'FINOLA' contig containing *CsFT1a* and *b* (a, QKVJ02000894.1|:189,639-200,768) was used as a query in a BLAST search against the *C. sativa* reference genome ('CBDRx', GCF_900626175.2) revealing multiple hits with high similarity to a 1 kb section of the large intron of *CsFT1a* (QKVJ02000894.1|:191,793-192,866). *CsFT1* of the reference genome (arrowhead, NC_044379.1|:64,421,405-64,426,176) is included in the alignment as well. A consensus logo generated from the alignment in (b) revealed sequence signatures characteristic for helitron transposons (green background), which typically insert into AT sites (grey background) (c)(Xiong et al., 2014).
